# Deep learning guided design of dynamic proteins

**DOI:** 10.1101/2024.07.17.603962

**Authors:** Amy B. Guo, Deniz Akpinaroglu, Mark J.S. Kelly, Tanja Kortemme

## Abstract

Deep learning has greatly advanced design of highly stable static protein structures, but the controlled conformational dynamics that are hallmarks of natural switch-like signaling proteins have remained inaccessible to *de novo* design. Here, we describe a general deep-learning-guided approach for *de novo* design of dynamic changes between intra-domain geometries of proteins, similar to switch mechanisms prevalent in nature, with atom-level precision. We solve 4 structures validating the designed conformations, show microsecond transitions between them, and demonstrate that the conformational landscape can be modulated by orthosteric ligands and allosteric mutations. Physics-based simulations are in remarkable agreement with deep-learning predictions and experimental data, reveal distinct state-dependent residue interaction networks, and predict mutations that tune the designed conformational landscape. Our approach demonstrates that new modes of motion can now be realized through *de novo* design and provides a framework for constructing biology-inspired, tunable and controllable protein signaling behavior *de novo*.

## Main Text

The ability of proteins to switch between multiple conformations is important for many biological functions including molecular recognition, signaling, and enzyme catalysis (*1*). The populations of conformational states and their interconversions are governed by a protein’s energy landscape (*2*) - the relative free energies of conformational states and the barriers between them. Specific conformational changes and their dynamics are also important for regulation: Binding partners, environmental conditions, post-translational modifications, and allosteric effectors can modulate the free energy landscape and consequently protein function by changing the populations of active and inactive conformations. Protein conformational landscapes and functions have also been tuned by computationally predicted mutations (*3, 4*). Notably, many of the conformational changes important for biological functions do not involve large-scale structural rearrangements; rather, critical motions often consist of localized rotation, tilting, and sliding of secondary structure elements within a domain. These types of motion are hallmarks of central biological processes, such as the regulation of kinases (*5*) and signaling via G-protein coupled receptors (GPCRs) (*6*) (Fig. S1).

Despite the functional importance of intra-domain conformational changes in natural proteins, predictive design of similar dynamics *de novo* has remained elusive (*7*). One key challenge is the difficulty in parameterizing sufficiently accurate physics-based energy functions or learned sequence-structure models to design sequences that can adopt multiple conformations (multi-state design, (*8*)). The energetic differences between individual conformational states are typically subtle (as small as a few kJ/mol) (*9*). Moreover, the small energy gap between designs that are interconverting between defined folded states, versus adopting disordered states, further complicates *in silico* design. A second challenge is to generate alternative states *de novo* (not borrowed from existing proteins) in the first place. These states also have to be designable, i.e. there must exist an amino acid sequence that can adopt these alternative conformations.

Consequently, there are only a few examples of *de novo* designed dynamic proteins (*10*), and none on the scale of intra-domain geometry changes prevalent in natural regulators. Pioneering studies include the design of fold switches (*11, 12*), side chain dynamics (*13*), and controllable changes in coiled-coil assemblies (*14*). Most commonly, design of protein switches has involved domain-level rearrangements or hinge-like motions of rigid bodies, where conformational changes are driven by intermolecular protein-protein interactions and the multi-state design problem is simplified as either most atomic interactions within the rigid bodies remain the same (*15*) or one of the states becomes disordered (*16*). Despite this long-standing interest, there is no generalizable method to engineer dynamic proteins (*10*). As a result, most classes of conformational switch mechanisms, effectors, and their combinations have been inaccessible and hence completely unexplored by *de novo* design.

In this study, we test whether recent advances in deep learning for sequence design (*17, 18*) and structure prediction (*19*) can be integrated with physics-based methods to design non-native local and controllable conformational dynamics in proteins *de novo*, inspired by the scale and modes of conformational changes prevalent in natural signaling proteins. Specifically, we develop and experimentally validate a generalizable method to design protein switches that interconvert between multiple conformational states, which differ in the intra-domain relative geometry of secondary structure elements. As in biological regulation, we show that the designed conformational states can be modulated by both ligand concentration and allosteric perturbations. Finally, we show that physics-based analyses suggest a molecular mechanism for modulating these *de novo* designed conformational landscapes. Taken together, our approach makes new modes of motions accessible to *de novo* design, providing a basis for designing tunable and complex signaling behavior beyond what exists in nature (*7*).

### Design approach for dynamic proteins with tunable two-state equilibria

We sought to develop a generalizable approach to design sequences with multiple energetic minima in structure space, each corresponding to a well-defined conformational state (Fig. 1A). Specifically, we focused on conformational changes involving intra-domain reorientation of secondary structural elements, mimicking a dominant mechanism in naturally occurring signaling proteins (in contrast to the easier problem of movement of otherwise static sub-domains about a hinge). We also aimed to design mechanisms for modulating the conformational equilibrium by inputs such as orthosteric ligands (binding within the region of conformational change) and allosteric perturbations (acting at distal sites).

**Fig. 1.**
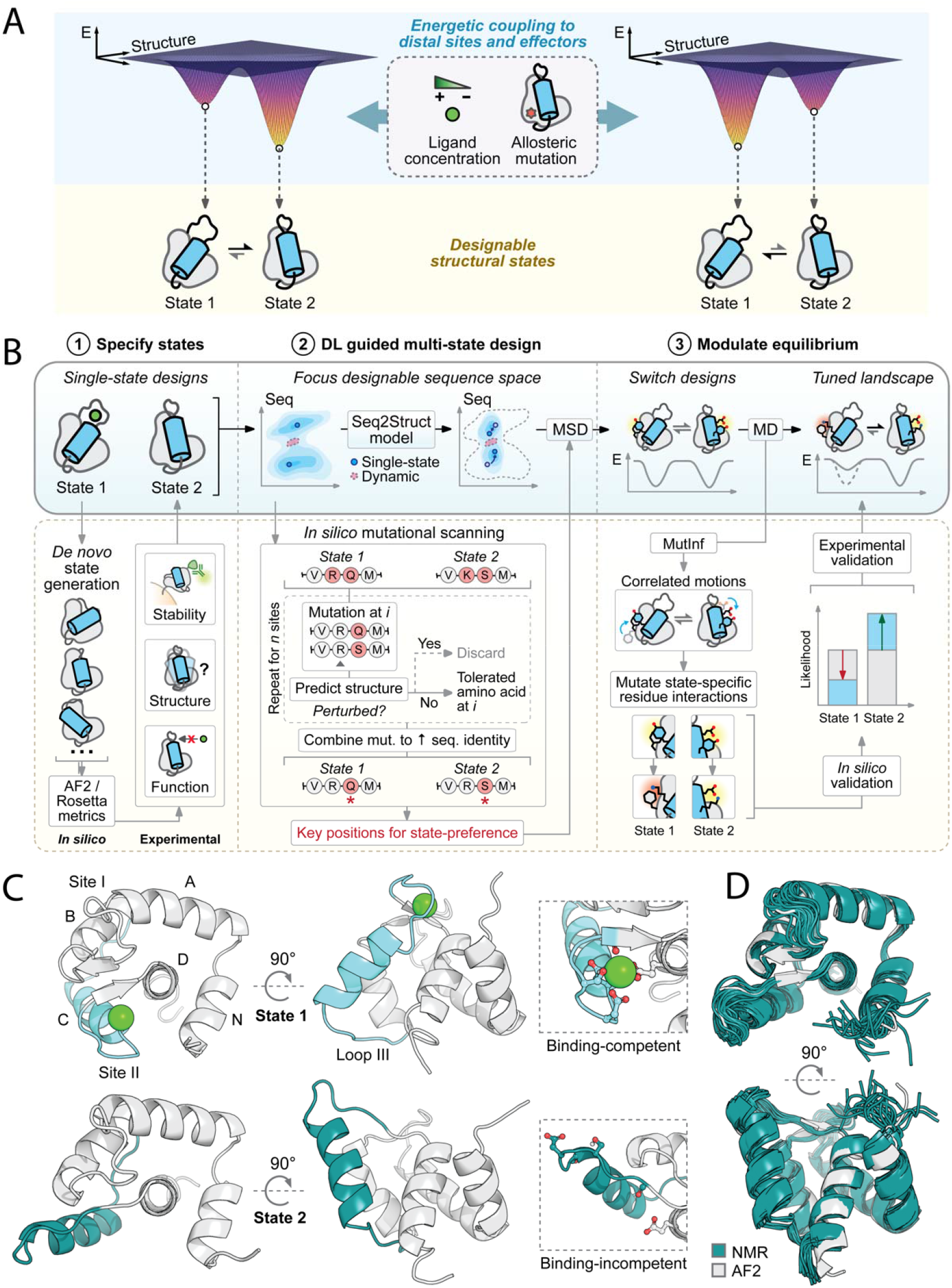
Generalizable approach for the deep learning guided design of dynamic proteins. (**A**)Schematic of design goal to engineer dynamic proteins in a two-state equilibrium that can be controlled by orthosteric ligands and allosteric perturbations. (**B**) Main stages of the approach: (1)*De novo* generation of alternative states that differ in their geometry (light blue region) using systematic conformational sampling (Fig. S2), followed by *in silico* and experimental validation of single state designs. (2) Deep learning (DL) guided sequence/structure search to focus sampling during multi-state design (MSD) at key positions for determining state preference and their neighbors. (3) Combination of physics-based molecular dynamics (MD) simulations, mutual information analysis (MutInf), and deep learning models to determine state-specific residue interaction networks and identify mutations capable of modulating the conformational landscape, followed by experimental validation. (**C-D**) Application to generate two designable states with distinct conformations coupled to ligand binding (Ca^2+^). (C) (Top row, light blue) Binding-competent state 1 structure with Ca^2+^ binding site (inset) (PDB ID: 1SMG) shown in two orientations. (Bottom row, teal) *De novo* generated alternative (binding-incompetent) state 2 model shown in two orientations. To couple Ca^2+^ binding to the designed conformational change, the Ca^2+^ binding site is significantly reshaped in state 2 to disfavor Ca^2+^ binding (inset). (D) Overlay of the NMR structure of a state 2 single-state design (teal) with its AF2 prediction (grey) shows excellent agreement (Cα RMSD = 0.98Å, excluding loops).

Our design approach (Fig. 1B) involves three general stages: The first stage specifies alternative structural states through (i) generation of a library of *de novo* candidate states that differ in their local geometries using systematic physics-based conformational sampling (*20*) (Fig. S2, Methods), and (ii) assessment of the designability of these states. To assess single-state designability of the generated backbones, we designed sequences for each state, evaluated them *in silico* (Methods), and characterized them experimentally. The second stage performs a deep learning guided search in sequence and structure space that (i) restricts the search space by identifying the minimal set of residues required to define each state, and (ii) designs sequences at these positions that are simultaneously compatible with pairs of states (multi-state design). The third stage seeks to identify perturbations that modulate the designed protein conformational landscape by (i) integrating physics-based simulations and deep learning predictions to determine state-specific interaction networks and (ii) predicting mutations leading to tunable conformational switching.

For the proof-of-concept application described here, we required the conformational landscape to be responsive to a ligand input, where the ligand-binding site (orthosteric site) changes conformations between states and thereby couples ligand binding to conformational switching.

For simplicity, we used an engineered Ca^2+^ binding protein (derived from the N-terminal domain of troponin C) as our ligand-binding-competent starting state (“state 1”) (Fig. 1C, top row). The wild-type protein consists of two EF hand motifs (sites I and II), which both bind Ca^2+^ in the low micromolar range. We instead used a variant (PDB ID: 1SMG) with an E41A point mutation in site I, which weakens the affinity of site I to the millimolar range while retaining micromolar affinity in site II (*21*). Additionally, the E41A mutant does not undergo conformational change upon Ca^2+^ binding (Fig. S3). By using an existing binding state as a starting point, we demonstrate here how our protocol can be generally applied to engineering controllable conformational changes into natural proteins; alternatively, one could generate a (static) binding-competent state *de novo* (*22*).

### Generation and experimental validation of de novo designed alternative states

To generate a structurally defined alternative state (“state 2”) (Fig. 1C, bottom row), we sampled *de novo* orientations of a contiguous protein segment including loop III, helix C, and Ca^2+^ binding site II (which we refer to as the “reshaped region”) (Fig. S2) using the loop-helix-loop unit combinatorial sampling algorithm (LUCS), previously shown to be capable of generating static proteins that differ in the local geometry of protein segments (*20*) (Methods). We only kept design models with the same length as our input but allowed the secondary structure in the reshaped region to vary (i.e., some regions may be part of a loop in the input but adopt a helical structure in the output or vice versa, a common transition in natural proteins). We then used Rosetta to design sequences optimal for each of the output models (“single-state designs”) and filtered these designs computationally (Methods). This procedure generated a library of approximately 1×10^3^ diverse conformations with an average Cα RMSD of 7.1Å in the reshaped region, from which we selected 11 designs (each corresponding to a unique backbone) for experimental testing.

To rapidly screen the *de novo* state 2 sequences experimentally, we displayed each design fused to a C-terminal c-Myc tag on the surface of yeast (Methods) and used surface display levels (which are known to be correlated with stability) as a proxy for designability. Although we only tested a few single-state designs in this study, one could screen thousands of designs with yeast display to identity many more designable conformational states. 10 out of 11 designs (all except #615) showed high surface display levels (Fig. S4). We decided to further characterize design #6306 as it had a significantly different conformation in the reshaped region compared to state 1 (Fig. 1C, bottom row), involving both rotation and translation of the reshaped helix C. Moreover, the Ca^2+^ binding loop was significantly restructured and partially helical, resulting in an unfavorable conformation for binding. We solved the nuclear magnetic resonance (NMR) structure of #6306 and saw excellent agreement (Cα RMSD = 0.98Å, excluding loops) between the experimentally solved structure (teal) and the Alphafold2 (AF2) model of the design (grey) (Fig. 1D, Fig. S5). In contrast to state 1 (1SMG), design #6306 did not show significant Ca^2+^ binding in site II at Ca^2+^ concentrations up to 1mM (Fig. S6), as expected due to significant restructuring of the binding site. These results confirm that the backbone of design #6306 is both designable and unfavorable for ligand binding, making it suitable as our binding-incompetent conformation (state 2) for two-state design.

### Multi-state design of dynamic proteins

We next aimed to identify sequences that were simultaneously compatible with the conformations of both state 1 and state 2. Rather than allowing all residues in the reshaped region and their contacting residue positions to be designable, we sought to generate multi-state designs able to populate both states despite having high sequence similarity. To do so, we used deep learning-based structure predictions (AF2) to shrink the searchable sequence space and focus sampling at key positions for determining state preference. We reasoned that this approach would allow us to better interpret sequence differences predicted to shift the preference for one state versus the other, identify potential sites for allosteric mutations, and facilitate experimental characterization. To achieve this, we used AF2 to identify mutations to design #6306 (predicted to adopt state 2) that would increase sequence similarity to 1SMG (state 1) without affecting the predicted structure (Methods) (Fig. 1B). We then used the resulting sequence with the highest sequence identity to state 1, which was still predicted to fold into state 2, as input for multi-state design (position-tied ProteinMPNN) (Methods, Fig. S7A-B) (*17*). Sites where mutations caused significant structural perturbations were typically positions where there were large changes to solvent accessible sidechain surface area, hydrogen bonding networks, or steric packing between states (Table S1). The final set of multi-state designable residues included these positions and their neighbors, decreasing the set of designable residues from 37 to 25 (Table S2). The structures of putative switch design sequences were then predicted with AF2 to assess their compatibility with both states.

Our design approach identified a family of sequences that had AF2 structure predictions that were either in state 1, state 2, or a combination of both including structural intermediates (Fig. 2A). The designs differed from the original state 1 sequence (1SMG) by n=18 mutations and from the high sequence identity single-state state 2 design by n=15 mutations, but strikingly differed from each other at only one residue position, 89, which was located outside the reshaped region and distal from the Ca^2+^ binding site. Position 89 was hence predicted to act as an allosteric site modulating state populations. In particular, smaller hydrogen bond donors and acceptors at position 89 were biased toward state 2 by forming a hydrogen bond with the backbone of loop III, bringing it closer to the central helix D. Conversely, bulky and/or hydrophobic amino acids pushed loop III outward into a conformation more consistent with state

**Fig. 2.**
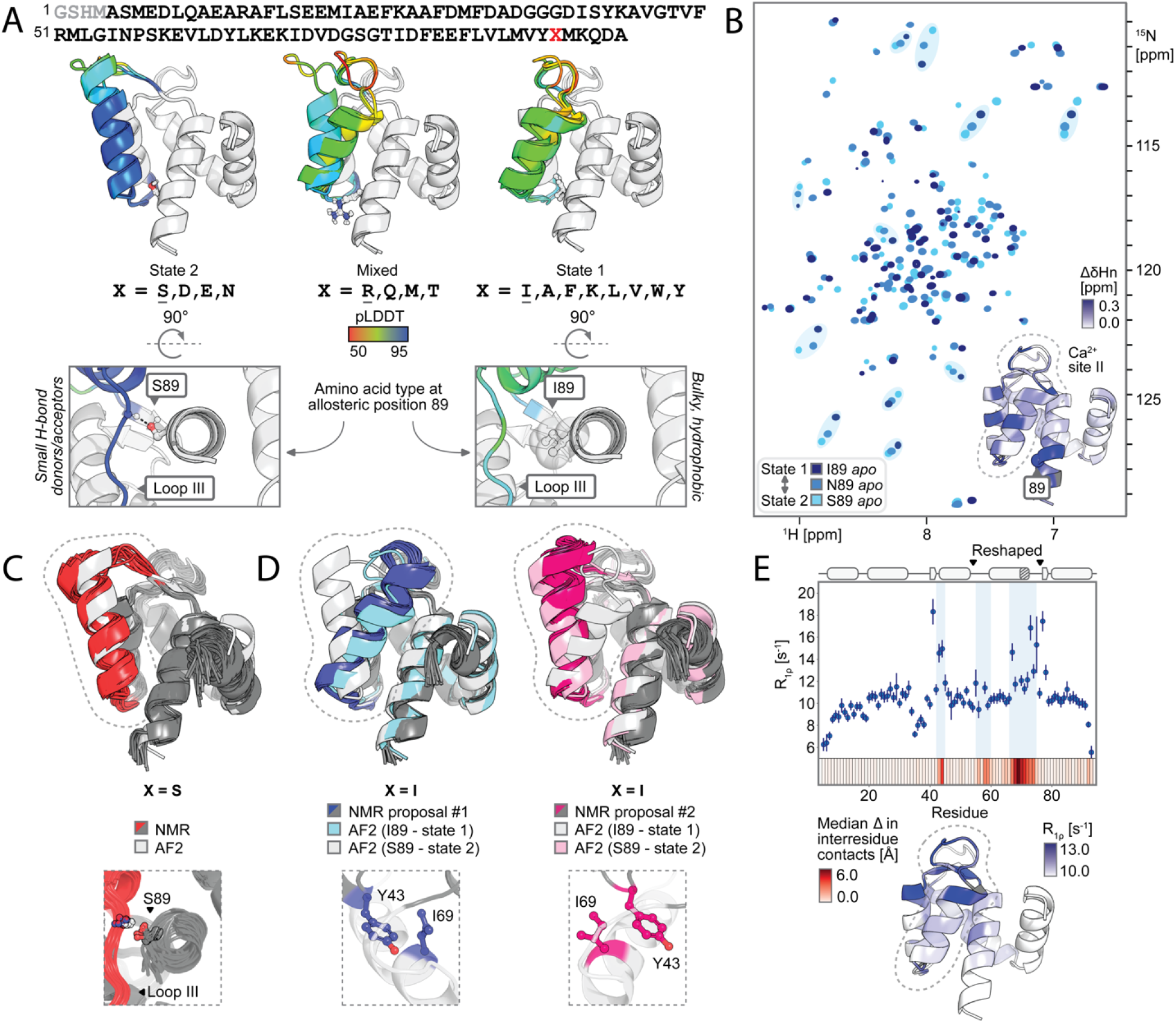
Two-state equilibrium in fast exchange shifted by allosteric mutations. (**A**) The protocol in Fig. 1B predicted a family of sequences differing only at position 89 (X), where the amino acid identity at position X determined whether the 5 AF2 predicted models were entirely in state 2 (left), mixed (middle), or entirely in state 1 (right). Depicted AF2 models (grey cartoons, with reshaped region colored by AF2 pLDDT) are for the underlined amino acid at position X (shown as sticks). Small polar residues favored state 2 by hydrogen-bonding with the backbone of loop III (bottom, left), while bulky and/or hydrophobic residues favored state 1 by pushing loop III away from the central helix (bottom, right). (**B**) ^1^H,^15^N-HSQC spectra of S89, N89, and I89, with several well-resolved peaks (shaded ovals) showing chemical shift changes consistent with a two-state equilibrium in fast exchange between state 1 preferred (I89), state 2 preferred (S89) and intermediate (N89). Inset shows ^1^Hn chemical shift changes between I89 and S89 colored on the AF2 model of I89, consistent with the designed conformational change in the reshaped region (the reshaped region is circled by dashed line in panels B-E). (**C**) Agreement between the NMR structure of S89 (red) and its AF2 prediction (grey) (Cα RMSD = 1.31Å excluding loops). Inset shows the hydrogen bond formed between S89 and loop III in the NMR structure consistent with AF2 predictions in (A). (**D**) NMR models for I89 were consistent with sampling both state 1 (blue, left) (Cα RMSD = 1.67Å excluding loops) and state 2 (pink, right) (Cα RMSD = 1.31Å excluding loops), with key residue ILE 69 buried in proposal #1 (state 1) and solvent-exposed in proposal #2 (state 2). (**E**) ^15^N near-resonance R_1ρ_ relaxation rates for design I89 plotted per residue (top) and visualized on the AF2 structure of design I89 (bottom) indicate low microsecond exchange in the regions predicted to undergo significant conformational changes (median predicted change in Cβ-Cβ distances between states shown on the x axis colored by magnitude). Residue numbering of all designs includes the N-terminal thrombin cleavage site scar (grey in (A)).

1. Moreover, the AF2 confidence metric (pLDDT) of the reshaped region distal to site 89 for this sequence family varied considerably depending on the amino acid identity at residues 89 (Fig. S7C).

### Allosteric modulation of the conformational landscape by mutation

To test the prediction that our designs will adopt two different defined conformations whose relative populations are dependent on the amino acid identify at position 89, we selected five designs that covered a range of AF2 predicted behaviors: state 2 preferred (S89, N89), state 1 preferred (I89, K89), and mixed (R89). We recorded ^1^H,^15^N-HSQC NMR spectra for each of these designs (Fig. 2B, Fig. S8) and focused structural characterization on three representative designs (S89, N89, I89). Remarkably, these spectra were drastically different even though the designs differed only by a single point mutation: 67 out of the 92 backbone amide peaks in the spectrum of I89 (state 1 preferred) had significantly different chemical shifts (ΔδHn > 0.03ppm or ΔδN > 0.4ppm) compared to that of S89 (state 2 preferred), suggesting different chemical environments of these residues (Fig. 2B). Moreover, for well-separated peaks, the chemical shifts of S89 and I89 were at the two ends of a range, with N89 being intermediate. This finding is consistent with the designed proteins being in equilibrium between two states in fast exchange on the NMR timescale where the observed chemical shifts reflect the population average of the two states. To analyze this behavior further, we assigned the backbone ^1^Hn and ^15^N chemical shifts of S89 and I89 and plotted the differences on the AF2 model of I89 (Fig. 2B, Table S3). We found that the changes to chemical shifts were not merely localized near the mutated position 89, but also at more distal residues throughout the reshaped region, including Ca^2+^ binding site II and its neighboring residues, consistent with a change in the ensemble averaged conformation of the reshaped region.

To assess whether our designs indeed adopted the two specified designed conformational states, we solved the structures for S89 and I89 by NMR (Fig. 2C-D). The structure of S89 was in excellent agreement with its AF2 prediction (state 2, Cα RMSD = 1.31Å, excluding loops), including in core hydrophobic residues (Fig. S5), consistent with the high average pLDDT (high confidence) of that design. In contrast, for I89, automated structure determination using ARTINA (Methods) did not converge to a single proposed conformation in the reshaped region. This observation is in line with the lower average pLDDT of I89 in the reshaped region compared to S89 (Fig. S7C). Instead, the top two structures proposed by ARTINA resembled the designed state 1 and state 2 conformations, respectively (Fig. 2D, Fig. S5), based on assigned distance restraints consistent with both states 1 and 2 (Fig. S9). The residue predicted to undergo the largest conformational change in our computational models (ILE 69) was buried in proposal #1 but solvent-exposed in proposal #2, hallmarks of the state 1 and 2 conformations, respectively. Taken together, our structures confirm that the designs adopt the two intended conformations but with different populations. While S89 primarily adopts state 2 as predicted, I89 samples both state 1 and state 2, and a single-state structure cannot fully account for all ensemble-averaged distance restraints simultaneously.

To probe the suggested dynamics in design I89 more directly, we first collected a series of ^1^H,^15^N-HSQC spectra from 5°C to 35°C at 5°C intervals. We observed temperature-dependent changes in peak intensity localized to residues in the reshaped region and their neighbors, consistent with changes in chemical environment due to dynamics in the reshaped region (Fig. S10). Moreover, peak intensities were higher at 35°C compared to 5°C, as expected for a system being in fast exchange at higher temperatures and slowing to intermediate exchange with decreasing temperature, manifesting as line broadening. Additionally, we measured R_1ρ_ values for design I89 with rotating frame relaxation dispersion. We observed higher R_1ρ_ values for residues in the reshaped region and their neighbors, again consistent with the reshaped region being in a two-state equilibrium as intended and exchanging on the low microsecond timescale (Fig. 2E, Table S4).

### Orthosteric modulation of the conformational landscape by ligand binding

To probe whether Ca^2+^ binding also modulates the state populations by preferentially stabilizing state 1, we recorded ^1^H,^15^N-HSQC spectra with and without Ca^2+^ for each point mutant (Fig. S11). As expected from the designed conformational change, we found that Ca^2+^ addition caused significant chemical shift perturbations (ΔδHn > 0.03ppm or ΔδN > 0.4ppm) throughout the reshaped region and its neighboring residues, affecting approximately 30 additional peaks when compared to the single-state binding-incompetent design #6306 (Fig. 3A, Table S5). Moreover, given that AF2 predicted primarily the binding-incompetent structure (state 2) for S89 and N89 and primarily the binding-competent structure (state 1) for I89 and K89 (Fig. 2A), the direction of chemical shift changes with Ca^2+^ was indeed consistent with a shift in the equilibrium toward the binding-competent state 1 (Fig. 3B). Notably, S89 and N89 (state 2 preferred) had perturbations that were smaller in magnitude compared to K89 and I89 (state 1 preferred). R89, which had a bimodal AF2 prediction (Fig. 2A), had approximately 40 minor peaks in the absence of Ca^2+^, consistent with slow exchange between two states. For this design, addition of Ca^2+^ lowered the intensity of many peaks, indicating a shift in exchange regime from slow to intermediate. Finally, we solved the NMR structure of I89 in the presence of Ca^2+^. The *holo* structure was in excellent agreement with our computational state 1 model (Cα RMSD = 1.34Å, excluding loops) and the backbone conformation of binding site II was consistent with an EF hand binding motif (Fig. 3C). Though *holo* I89 had many more distance restraints consistent with state 1 and less consistent with state 2 compared to *apo* I89, we still observed several nOe-derived distance restraints consistent with state 2 even in the presence of excess Ca^2+^, suggesting residual dynamics (Fig. S12). Taken together, these results suggest that the tested family of sequences adopted the two designed conformational states of the reshaped region in solution, where the populations of these states can be modulated by both allosteric mutations (Fig. 2) and Ca^2+^ binding (Fig. 3).

**Fig. 3:**
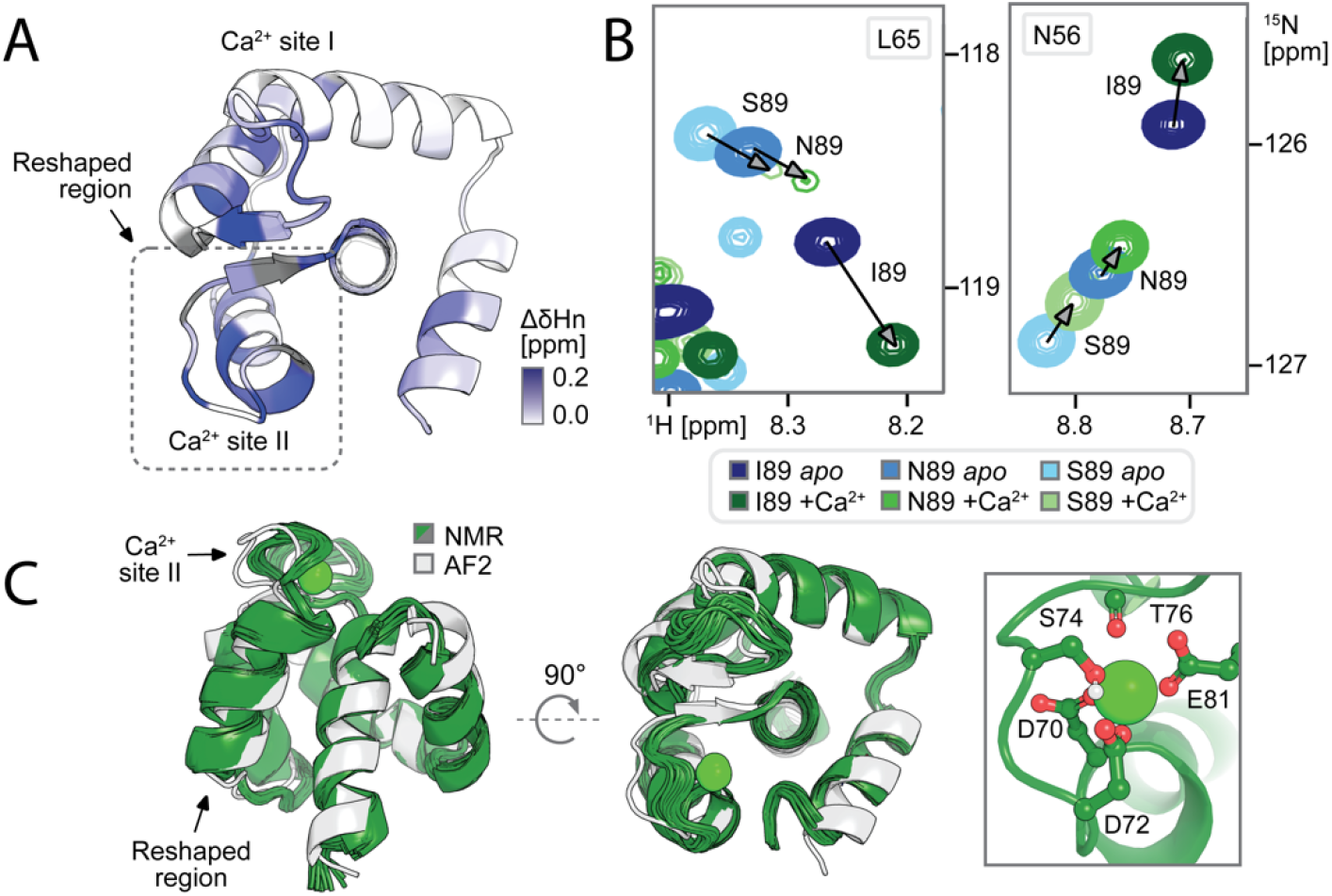
Modulation of the conformational landscape by ligand binding. (**A**) Changes in I89 ^1^Hn chemical shifts upon adding 10eq of Ca^2+^ visualized on the AF2 model, showing significant chemical shift perturbations distal to the Ca^2+^ binding sites, particularly in the reshaped region and in neighboring residues. (**B**) ^1^H,^15^N-HSQC spectra of S89, N89, and I89 with 10eq of Ca^2+^ show ligand-dependent chemical shift changes for residues in the reshaped region consistent with Ca^2+^ shifting the population distribution toward state 1 (arrows). (**C**) The Ca^2+^ bound NMR structure of I89 is excellent agreement with its AF2 prediction (Cα RMSD = 1.34Å, excluding loops), and the observed binding site backbone is consistent with the known EF hand binding motif with modeled-in Ca^2+^ (right).

### Integration of physics-based simulations to tune conformational equilibria

To further probe the atom-level interactions underlying the designed conformational switching, we ran molecular dynamics (MD) simulations for designs I89 and S89 with and without Ca^2+^ (Fig. 4A). Strikingly, throughout the trajectory initiated from the AF2 model without Ca^2+^, design I89 switched between state 1 and state 2 in excellent agreement with our design (Fig. 4A, Fig. S13) with a timescale of exchange consistent with our R_1ρ_ data (Fig. 2E). In contrast, we did not observe conformational switching when simulating design I89 with Ca^2+^, and Ca^2+^ remained bound to site II during the simulation (Fig. 4A, inset), in agreement with the experimental data showing that Ca^2+^ stabilizes state 1. Design S89 remained in the state 2 conformation throughout the entire course of a 1μs simulation under both the Ca^2+^-free condition and in the presence of Ca^2+^ (Fig. S14); the latter could be explained by insufficient sampling, although the lack of observed dynamics on this timescale is consistent with our AF2 prediction (Fig. 2A), the higher (more confident) average AF2 pLDDT of S89 compared to I89 (Fig. S7C), the NMR structure (Fig. 2C), and the Ca^2+^ binding data (Fig. 3B). In summary, the MD trajectories provide compelling support of a two-state equilibrium between the designed states on a low microsecond timescale for design I89, show that Ca^2+^ preferentially stabilizes state 1 in I89, and are consistent with allosteric modulation of the designed switch at position 89.

**Fig. 4:**
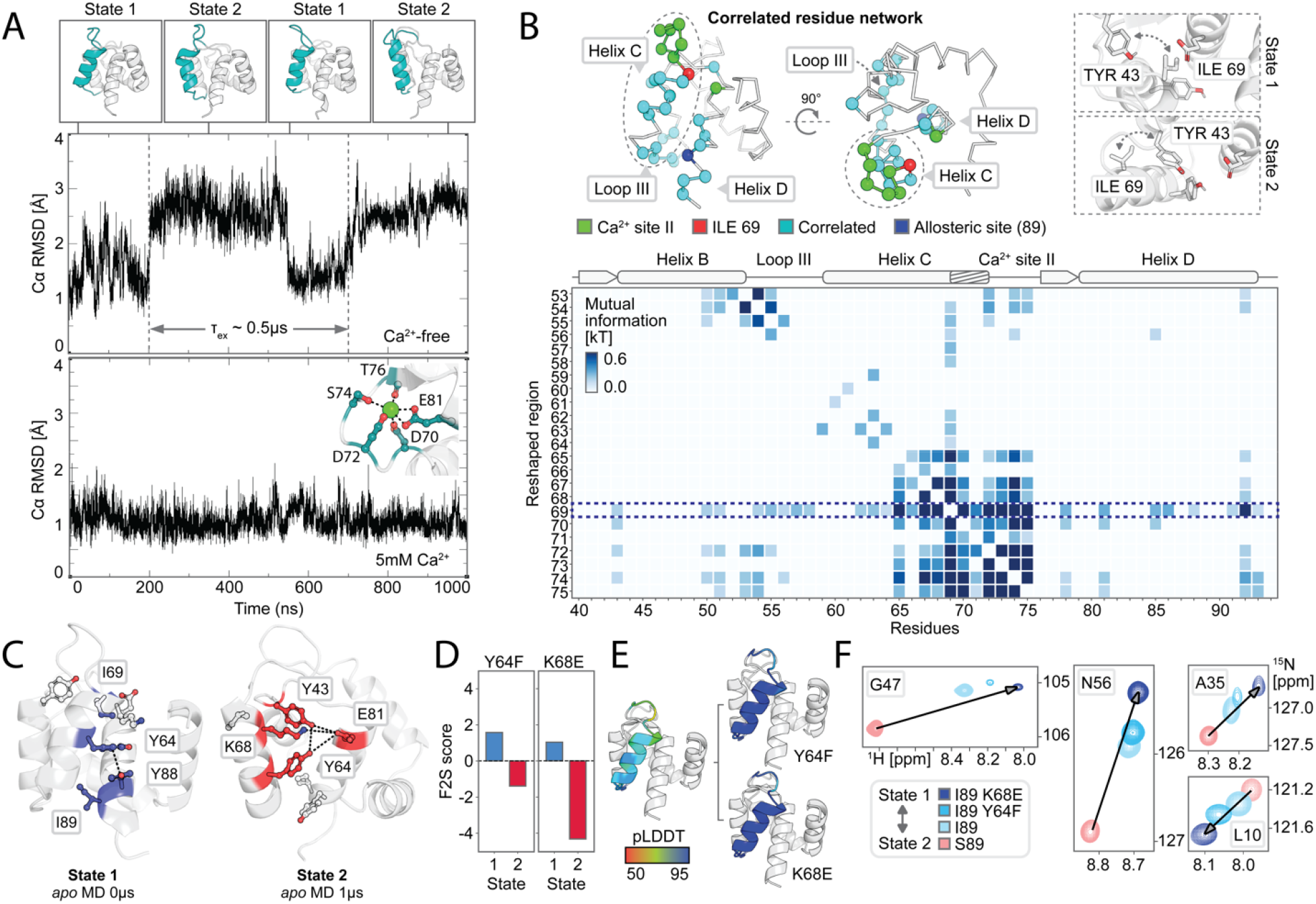
Physics-based simulations reveal molecular mechanisms underlying switch behavior. (**A**) Ca^2+^-free 1μs molecular dynamics (MD) trajectory of I89 showing the expected transitions between a binding-competent state 1 and binding-incompetent state 2 conformation (cyan helix, top) with a timescale of exchange of approximately 0.5μs. The protein remains in state 1 in the presence of Ca^2+^ (bottom). C⍰ RMSD is measured for the reshaped helix compared to the starting coordinates and coordinates Ca^2+^ consistent with an EF hand (inset). (**B**) Mutual informational analysis (heatmap) revealing a correlated network of residues (light blue) connecting the Ca^2+^ binding loop (green) with the allosteric mutation site (dark blue), shown in two orientations (top panels, reshaped helix circled by a dashed line). Panels on the right show interaction details of a key residue, ILE 69 (red, highlighted by dashed box in heatmap), linking the two regions. ILE 69 becomes solvent exposed in state 2, enabling the formation of distinct contacts by TYR 43. (**C**) State-specific interactions for state 1 and state 2 (colored) observed during a 1μs Ca^2+^-free MD simulation for I89. (**D-F**) In silico and experimental validation of state-specific interaction networks using mutations. (D) Frame2Seq (F2S) predictions for Y64F and K68E; score refers to the negative log-likelihood difference between the mutated and original sequence, where the negative change in score predicts the mutations disfavor state 2. (E) AF2 predictions showing higher pLDDT (greater confidence) for state 1 for mutants on the right compared to the original sequence on the left. (F) ^1^H,^15^N-HSQC NMR data showing chemical shift changes consistent with an increased state 1 population in the mutants (arrows denoting direction of shifts from state 2 (S89) towards a larger population of state 1 (I89)).

Because of the remarkable agreement between the design predictions (Fig. 2A), MD simulations (Fig. 4A), and NMR data (Fig. 2B-E; Fig. 3), we next asked whether the MD results could (i) explain the mechanism of allosteric modulation and (ii) make testable prospective predictions to further validate this mechanism. We first performed a mutual information analysis of side chain torsional dynamics (*23*) in the *apo* I89 MD trajectory. We observed a network of hydrophobic core residues coupling torsional motions of the Ca^2+^ binding site II (residues 70-76) to distal residues in loop III (residues 50-58) and helix D (residues 80-94) (Fig. 4B). Allosteric residue 89 directly faces loop III from central helix D. Combined with our experimental results on allosteric modulation (Fig. 2), our MD results therefore suggest a mechanism where the amino acid identity at residue 89 causes differences in sterics (I89) or hydrogen bonding (S89) interactions between helix D and loop III (Fig. 2A), which in turn couple through the identified correlated hydrophobic network to the Ca^2+^ binding site and reshaped region (Fig. 4B). Further analysis of the mutual information data revealed two unique interaction networks that are specific for state 1 or state 2, respectively (Fig. 4C), and involved motions of ILE 69 as a key facilitator of communication between distal regions. State 1 appeared to be stabilized by hydrophobic interactions involving burial of ILE 69, π-π stacking between TYR 64 and TYR 88, and less steric bulk near ILE 89. Conversely, in state 2 ILE 69 is pointed outwards to the surface, allowing for TYR 43 to form new contacts in network of hydrogen-bonding and electrostatic interactions mediated by TYR 43, TYR 64, LYS 68, and GLU 81. Taken together, the mutual information analysis of the MD trajectory suggests two extensive state-specific interaction networks that link the reshaped region and the allosteric site 89.

To test these state-specific interactions, we predicted mutations of the identified networks that would favor state 1 by (i) scoring with a structure-conditioned masked language model (Frame2seq) (*18*) and (2) predicting the mutant structures with AF2. We reasoned that a Y64F mutation should be disruptive in state 2, as it is unable to hydrogen bond with TYR 43 and GLU 81, but neutral in state 1, where PHE can still form a π-π stacking interaction with TYR 88.

Likewise, a K68E mutation should also be disruptive in state 2, as it cannot form a stabilizing electrostatic interaction with GLU 81, but neutral in state 1 where it is solvent-exposed. In both cases, Frame2seq predicted a higher likelihood for the mutated amino acid compared to the original amino acid given state 1 and a lower likelihood given state 2, AF2 predicted these mutants to adopt state 1 with higher confidence compared to the original I89 sequence, and the ^1^H,^15^N-HSQC spectra were consistent with the expected shift in population toward state 1 (Fig. 4D-F). In summary, our results explain the observed allosteric modulation of the conformational landscape by mutations at the lynchpin position 89 and validate predictions to further tune the switch equilibrium.

## Discussion

Our results demonstrate a general approach to design proteins with two distinct conformational states specified by the designer whose interconversion can be modulated both by ligand concentration (orthosterically) and by mutations to distal sites (allosterically). The approach generalizes beyond prior *de novo* designed switches based primarily on domain replacement or hinge-like motions (*10*) where most of the atomic interactions between the rigid bodies remain constant (*15, 16*). In contrast, our designs interchange between distinct sets of atomic interactions and demonstrate that new modes of motions – inspired by those present in regulator superfamilies such as kinases and GPCRs - can now be realized through *de novo* design, greatly expanding accessible functional space.

One notable observation is the strong correspondence between the deep-learning based predictions, experimental data, and physics-based simulations. This agreement provided testable hypotheses on the mechanisms underlying the bistability of the switch and allowed us to modulate the conformational landscape at the atom-level. We attribute these findings at least in part to specific features of our approach, in particular the search through sequence and structure space that narrowed designable positions to those predicted to be key determinants of the engineered conformational change (Fig. 1B). The speed and inference offered by deep-learning-based protein sequence design and structure prediction enabled this approach that, ultimately, designed distinct residue networks stabilizing the two structural states. This level of insight into a designed system is essential to advance *de novo* design of allosteric regulation.

The synergy between deep learning and physics-based simulations - demonstrated here for the first time for the *de novo* design of dynamic proteins - may be useful for developing future design methods that would enable predictive control over the timescale of exchange (which we did not consider here). Emerging approaches to this problem include training models on simulation and/or experimental data (*24, 25*) and using machine learning to parameterize new force-fields (*26, 27*). Future development of deep learning integrated with unique dynamics data offered by experiment and simulation could allow for one-shot conformational ensemble prediction (*28, 29*) to design sequences given a user-defined conformational landscape. It is encouraging that our approach can already successfully design motions involving intricate changes to the inter-residue interaction network between states, despite the individual models for sequence and structure prediction never having been explicitly trained for such a task.

Functional conformational changes in natural proteins are often complex, typically not well-understood, and hence difficult to study and manipulate using engineering approaches. In contrast, our design method both generates two-state conformational changes similar to those in nature, as well as successful predictions how to modulate the populations of these states. To our knowledge the proof-of-concept application here is the first example of designing a *de novo* alternative state into a naturally occurring protein. This result suggests that our approach may be useful for controlling natural functions through tunable conformational switching, which would open many new avenues to study and manipulate the roles of natural switches in the cellular context. Moreover, our approach should be useful to generate entirely bio-orthogonal protein switches. The approach and concepts described here hence have broad relevance to both modulate and deconstruct the requirements for natural signaling and construct complex switch-like signaling systems entirely from the ground up.

## Acknowledgments

We thank members of the Kortemme lab for discussion and Philipp Huettemann, Dr. Stephanie Crilly, and Dr. Robert G. Alberstein for comments on the manuscript.

## Author contributions

A.B.G. and T.K. developed the conceptual approach for designing dynamic proteins. A.B.G. designed, screened, and characterized *de novo* state 2 designs.

A.B.G. performed multi-state design and conceptualized, designed, screened, and characterized allosteric point mutants. A.B.G. collected and processed NMR data with aid and supervision from M.J.S.K. A.B.G. ran MD simulations and mutual information analysis.

D.A. performed Frame2seq scoring. T.K. provided guidance and resources. A.B.G. and T.K. wrote the manuscript with contributions from the other authors. All authors read and commented on the manuscript.

## Supplementary Materials

Materials and Methods

Figs. S1 to S16

Tables S1 to S10

References (*30*–*46*)

## References and Notes

1. D. D. Boehr, R. Nussinov, P. E. Wright, The role of dynamic conformational ensembles in biomolecular recognition. Nature Chemical Biology 5, 789–796 (2009).

2. K. Henzler-Wildman, D. Kern, Dynamic personalities of proteins. Nature 450, 964–972 (2007).

3. K. Y. M. Chen, D. Keri, P. Barth, Computational design of G Protein-Coupled Receptor allosteric signal transductions. Nature Chemical Biology 16, 77–86 (2020).

4. A. D. St-Jacques, J. M. Rodriguez, M. G. Eason, S. M. Foster, S. T. Khan, A. M. Damry, N. K. Goto, M. C. Thompson, R. A. Chica, Computational remodeling of an enzyme conformational landscape for altered substrate selectivity. Nature Communications 14, 6058 (2023).

5. M. Huse, J. Kuriyan, The conformational plasticity of protein kinases. Cell 109, 275–282 (2002).

6. A. S. Hauser, A. J. Kooistra, C. Munk, F. M. Heydenreich, D. B. Veprintsev, M. Bouvier, M. M. Babu, D. E. Gloriam, GPCR activation mechanisms across classes and macro/microscales. Nature Structural and Molecular Biology 28, 879–888 (2021).

7. T. Kortemme, De novo protein design—From new structures to programmable functions. Cell 187, 526–544 (2024).

8. J. A. Davey, R. A. Chica, Multistate approaches in computational protein design. Protein Science 21, 1241–1252 (2012).

9. S. J. Fleishman, D. Baker, Role of the biomolecular energy gap in protein design, structure, and evolution. Cell 149, 262–273 (2012).

10. R. G. Alberstein, A. B. Guo, T. Kortemme, Design principles of protein switches. Current Opinion in Structural Biology 72, 71–78 (2022).

11. X. I. Ambroggio, B. Kuhlman, Computational design of a single amino acid sequence that can switch between two distinct protein folds. Journal of the American Chemical Society 128, 1154–1161 (2006).

12. B. Ruan, Y. He, Y. Chen, E. J. Choi, Y. Chen, D. Motabar, T. Soloman, R. Simmerman, T. Kauffman, D. T. Gallagher, J. Orban, P. N. Bryan, Design and characterization of a protein fold switching network. Nature Communications 14, 431 (2023).

13. J. A. Davey, A. M. Damry, N. K. Goto, R. A. Chica, Rational design of proteins that exchange on functional timescales. Nature Chemical Biology 13, 1280–1285 (2017).

14. J. A. Cross, W. M. Dawson, S. R. Shukla, J. F. Weijman, J. Mantell, M. P. Dodding, D. N. Woolfson, A de novo designed coiled coil-based switch regulates the microtubule motor kinesin-1. Nature Chemical Biology 20, 916–923 (2024).

15. F. Praetorius, P. J. Y. Leung, M. H. Tessmer, A. Broerman, C. Demakis, A. F. Dishman, A. Pillai, A. Idris, D. Juergens, J. Dauparas, X. Li, P. M. Levine, M. Lamb, R. K. Ballard, S. R. Gerben, H. Nguyen, A. Kang, B. Sankaran, A. K. Bera, B. F. Volkman, J. Nivala, S. Stoll, D. Baker, Design of stimulus-responsive two-state hinge proteins. Science 381, 754–760 (2023).

16. R. A. Langan, S. E. Boyken, A. H. Ng, J. A. Samson, G. Dods, A. M. Westbrook, T. H. Nguyen, M. J. Lajoie, Z. Chen, S. Berger, V. K. Mulligan, J. E. Dueber, W. R. P. Novak, H. El-Samad, D. Baker, De novo design of bioactive protein switches. Nature 572, 205–210 (2019).

17. J. Dauparas, I. Anishchenko, N. Bennett, H. Bai, R. J. Ragotte, L. F. Milles, B. I. M. Wicky Courbet, R. J. de Haas, N. Bethel, P. J. Y. Leung, T. F. Huddy, S. Pellock, D. Tischer, F. Chan, B. Koepnick, H. Nguyen, A. Kang, B. Sankaran, A. K. Bera, N. P. King, D. Baker, Robust deep learning–based protein sequence design using ProteinMPNN. Science 378, 49– 56 (2022).

18. D. Akpinaroglu, K. Seki, A. Guo, E. Zhu, M. J. S. Kelly, T. Kortemme, Structure-conditioned masked language models for protein sequence design generalize beyond the native sequence space. bioRxiv 2023.12.15.571823 [Preprint] (2023). 10.1101/2023.12.15.571823.

19. J. Jumper, R. Evans, A. Pritzel, T. Green, M. Figurnov, O. Ronneberger, K. Tunyasuvunakool, R. Bates, A. Žídek, A. Potapenko, A. Bridgland, C. Meyer, S. A. A. Kohl, J. Ballard, A. Cowie, B. Romera-Paredes, S. Nikolov, R. Jain, J. Adler, T. Back, S. Petersen, D. Reiman, E. Clancy, M. Zielinski, M. Steinegger, M. Pacholska, T. Berghammer, S. Bodenstein, D. Silver, O. Vinyals, A. W. Senior, K. Kavukcuoglu, P. Kohli, D. Hassabis, Highly accurate protein structure prediction with AlphaFold. Nature 596, 583–589 (2021).

20. X. Pan, M. C. Thompson, Y. Zhang, L. Liu, J. S. Fraser, M. J. S. Kelly, T. Kortemme, Expanding the space of protein geometries by computational design of de novo fold families. Science 369, 1132–1136 (2020).

21. S. M. Gagné, M. X. Li, B. D. Sykes, Mechanism of direct coupling between binding and induced structural change in regulatory calcium binding proteins. Biochemistry 36, 4386– 4392 (1997).

22. R. Krishna, J. Wang, W. Ahern, P. Sturmfels, P. Venkatesh, I. Kalvet, G. R. Lee, F. S. Morey-Burrows, I. Anishchenko, I. R. Humphreys, R. McHugh, D. Vafeados, X. Li, G. A. Sutherland, A. Hitchcock, C. Neil Hunter, A. Kang, E. Brackenbrough, A. K. Bera, M. Baek, F. DiMaio, D. Baker, Generalized biomolecular modeling and design with RoseTTAFold All-Atom. Science 384, eadl2528 (2024).

23. C. L. McClendon, G. Friedland, D. L. Mobley, H. Amirkhani, M. P. Jacobson, Quantifying correlations between allosteric sites in thermodynamic ensembles. Journal of Chemical Theory and Computation 5, 2486–2502 (2009).

24. F. Noé, S. Olsson, J. Köhler, H. Wu, Boltzmann generators: Sampling equilibrium states of many-body systems with deep learning. Science 365, eaaw1147 (2019).

25. S. Mansoor, M. Baek, H. Park, G. R. Lee, D. Baker, Protein Ensemble Generation Through Variational Autoencoder Latent Space Sampling. Journal of Chemical Theory and Computation 20, 2689–2695 (2024).

26. A. E. P. Durumeric, N. E. Charron, C. Templeton, F. Musil, K. Bonneau, A. S. Pasos-Trejo, Y. Chen, A. Kelkar, F. Noé, C. Clementi, Machine learned coarse-grained protein force-fields: Are we there yet? Current Opinion in Structural Biology 79, 102533 (2023).

27. M. Majewski, A. Pérez, P. Thölke, S. Doerr, N. E. Charron, T. Giorgino, B. E. Husic, C. Clementi, F. Noé, G. De Fabritiis, Machine learning coarse-grained potentials of protein thermodynamics. Nature Communications 14, 5739 (2023).

28. G. Janson, G. Valdes-Garcia, L. Heo, M. Feig, Direct generation of protein conformational ensembles via machine learning. Nature Communications 14, 774 (2023).

29. S. Zheng, J. He, C. Liu, Y. Shi, Z. Lu, W. Feng, F. Ju, J. Wang, J. Zhu, Y. Min, H. Zhang, S. Tang, H. Hao, P. Jin, C. Chen, F. Noé, H. Liu, T.-Y. Liu, Predicting equilibrium distributions for molecular systems with deep learning. Nature Machine Intelligence 6, 558–567 (2024).

30. S. J. Fleishman, A. Leaver-Fay, J. E. Corn, E. Strauch, S. D. Khare, N. Koga, J. Ashworth, P. Murphy, F. Richter, G. Lemmon, J. Meiler, D. Baker, RosettaScripts: A scripting language interface to the Rosetta macromolecular modeling suite. PLoS One 6, e20161 (2020).

31. D. Gront, D. W. Kulp, R. M. Vernon, C. E. Strauss, D. Baker, Generalized fragment picking in Rosetta: design protocols and applications. PLoS One 6, e23294 (2011).

32. R. F. Alford, A. Leaver-Fay, J. R. Jeliazkov, M. J. O’Meara, F. P. DiMaio, H. Park, M. V. Shapovalov, P. D. Renfrew, V. K. Mulligan, K. Kappel, J. W. Labonte, M. S. Pacella, R. Bonneau, P. Bradley, R. L. Dunbrack, R. Das, D. Baker, B. Kuhlman, T. Kortemme, J. J. Gray, The Rosetta All-Atom Energy Function for Macromolecular Modeling and Design. Journal of Chemical Theory and Computation 13, 3031–3048 (2017).

33. R. D. Gietz, R. H. Schiestl, High-efficiency yeast transformation using the LiAc/SS carrier DNA/PEG method. Nature Protocols 2, 31–34 (2007).

34. E. V. Shusta, M. C. Kieke, E. Parke, D. M. Kranz, K. D. Wittrup, Yeast polypeptide fusion surface display levels predict thermal stability and soluble secretion efficiency. Journal of Molecular Biology 292, 949–956 (1999).

35. P. Klukowski, R. Riek, P. Güntert, Rapid protein assignments and structures from raw NMR spectra with the deep learning technique ARTINA. Nature Communications 13, 6151 (2022).

36. Y. Shen, F. Delaglio, G. Cornilescu, A. Bax, TALOS+: A hybrid method for predicting protein backbone torsion angles from NMR chemical shifts. Journal of Biomolecular NMR 44, 213–223 (2009).

37. S. P. Skinner, R. H. Fogh, W. Boucher, T. J. Ragan, L. G. Mureddu, G. W. Vuister, CcpNmr AnalysisAssign: a flexible platform for integrated NMR analysis. Journal of Biomolecular NMR 66, 111–124 (2016).

38. C. D. Schwieters, G. A. Bermejo, G. M. Clore, Xplor-NIH for molecular structure determination from NMR and other data sources. Protein Sci 27, 26–40 (2018).

39. H. Berman, K. Henrick, H. Nakamura, J. L. Markley, The worldwide Protein Data Bank (wwPDB): Ensuring a single, uniform archive of PDB data. Nucleic Acids Research 35, D301–D303 (2007).

40. N. C. J. Strynadka, M. N. G. James, Crystal structures of the helix-loop-helix calcium-binding proteins. Annual Review of Biochemistry 58, 951–998 (1989).

41. A. M. Sevy, T. M. Jacobs, J. E. Crowe, J. Meiler, Design of Protein Multi-specificity Using an Independent Sequence Search Reduces the Barrier to Low Energy Sequences. PLoS Computational Biology 11, e1004300 (2015).

42. P. Löffler, S. Schmitz, E. Hupfeld, R. Sterner, R. Merkl, Rosetta:MSF: a modular framework for multi-state computational protein design. PLoS Computational Biology 13, e1005600 (2017).

43. M. Mirdita, K. Schütze, Y. Moriwaki, L. Heo, S. Ovchinnikov, M. Steinegger, ColabFold: making protein folding accessible to all. Nature Methods 19, 679–682 (2022).

44. D. Van Der Spoel, E. Lindahl, B. Hess, G. Groenhof, A. E. Mark, H. J. C. Berendsen, GROMACS: Fast, flexible, and free. Journal of Computational Chemistry 26, 1701–1718 (2005).

45. P. Robustelli, S. Piana, D. E. Shaw, Developing a molecular dynamics force field for both folded and disordered protein states. Proceedings of the National Academy of Sciences of the United States of America 115, E4748–E4766 (2018).

46. J. Meier, R. Rao, R. Verkuil, J. Liu, T. Sercu, A. Rives, Language models enable zero-shot prediction of the effects of mutations on protein function. bioRxiv 2021.07.09.450648 [Preprint] (2021). 10.1101/2021.07.09.450648.

