## Supplementary Materials for "Deep learning guided design of dynamic proteins"

**The PDF file includes:**

Materials and Methods

Figs. S1 to S16

Tables S1 to S10

References

Materials and Methods

Residue numbering

Please note that all designs reported in this manuscript (including deposited structures/computational models, experimental data, and molecular dynamics simulation analyses) are indexed to include a four residue N-terminal thrombin cleavage scar (grey in Fig. 4A in the main manuscript). We adopted this residue numbering convention throughout (unless we are directly referring to the 1SMG PDB structure) to be consistent with the design constructs characterized structurally. We note that the scar was not explicitly modeled during generation of alternative *de novo* states, single-state design of these states, or multi-state design, which used a scar-less sequence with numbering starting at one (see Readme note for deposited scripts).

Generating alternative backbone conformations

The starting Ca^2+^ binding-competent structure (PDB ID: 1SMG) was obtained from the Protein Data Bank. Conformer 1 of the NMR ensemble was relaxed in Rosetta with all heavy atoms in the backbone and sidechains restrained.

relax.linuxgccrelease -database $ROSETTA/main/database -ex1 -ex2 -use_input_sc -flip_HNQ -no_optH false -relax:constrain_relax_to_start_coords -relax:coord_constrain_sidechains -relax:ramp_constraints false -s 1SMG.pdb

The loop-helix-loop unit (“reshaped region”, residues 49-72) involving the higher affinity Ca^2+^ binding loop (residues 68-72 and 77) was reshaped with loop-helix-loop unit combinatorial sampling (*20*) using loop libraries of length 2-5 residues. Note that these residue ranges follow 1SMG residue numbering, i.e. these residues correspond to residues 53-76 for the reshaped region and 70-76 and 81 for Ca^2+^ binding site II in our designs and presented data.

Only models having the same length as the input were written out. Since the input structure was primarily helical, alanine was used as the clash check residue rather than valine. Any outputs that had a reshaped helix orientation compared to the input (Cα RMSD < 3Å) were filtered out to generate alternative conformations that were experimentally distinguishable. Any outputs that had a reshaped helix Cα RMSD > 10Å compared to the input were also discarded, as these structures often lacked a well-packed hydrophobic core. This left n=921 candidate alternative *apo* backbone conformations for further analysis.

Single-state sequence design

A residue i was defined as pointing toward the reshaped region if the cosine of the angle between the vector pointing from Cα_i_ to Cα_j_ and the vector pointing from Cα_i_ to Cβ_i_ was greater than 0.5, where j is any residue in the reshaped region. Residues in the reshaped region as well as all residues within 10Å (Cα-Cα distance) from the reshaped region and pointing toward the reshaped region were designable, i.e. allowed to change amino acid identity and rotamer conformation. Residues within 8Å (Cα-Cα distance) and pointing toward designable residues were repackable, i.e. constant amino acid identity but allowed to change rotamer conformation. Residues in the reshaped region that were highly conserved in EF hand motifs were held fixed during design (i.e. D66, D68, G69, S70, G71, T72, E77, following PDB: 1SMG numbering convention). Allowable amino acids at designable residue positions were determined by the extent of residue burial using the Rosetta LayerDesign task operation (*30*). Cysteine and histidine were disallowed at all positions to avoid disulfide bond formation and pH sensitivity issues. Rosetta FastDesign and RotamerTrial protocols using extra rotamers for χ_1_ and χ_2_ (enabled by the ex1 and ex2 options in the ExtraRotamersGeneric task operation) were applied during design. Relevant scripts and command line prompts can be found at https://github.com/Kortemme-Lab/local_protein_sequence_design. 20 designs per backbone were generated.

*In silico* validation of single state designs

9-mer and 3-mer fragments for structure prediction were generated using the make_fragments.pl script distributed with Rosetta (*31*).

make_fragments.pl -verbose -id design_id -frag_sizes 3,9 -n_frags 200 -n_candidates 1000 sequence.fasta

A biased *ab initio* structure prediction simulation was run as a preliminary screen for the lowest scoring design (using the ref2015 Rosetta energy function (*32*)) of each backbone. This was done by using only the top three 9-mer/3-mer fragments closest in Cα RMSD to the desired backbone and generating 30 decoys. If low-RMSD decoys did not have low Rosetta energies or the lowest energy decoy was >3Å in Cα RMSD to the desired backbone, the design was removed from further analysis, as it would be unlikely to fold into the desired structure in an unbiased simulation. Full Rosetta *ab initio* structure prediction simulations (20,000 decoys, 200 fragments at each sliding window) were run for the remaining designs.

AbinitioRelax.linuxgccrelease -abinitio:relax -use_filters true -abinitio::increase_cycles 10 -abinitio::rg_reweight 0.5 -abinitio::rsd_wt_helix 0.5 -abinitio::rsd_wt_loop 0.5 -relax::fast -in:file:fasta sequence.fasta -in:file:frag3 <fragments_3mer_file> -in:file:frag9 <fragments_9mer_file> -psipred_ss2 <ss2_file_from_frag_generation> -nstruct <num_output> -out:sf <score_file_output> -out:file:silent <silent_file_output>

We selected 11 designs for experimental characterization (Fig. S4, yeast display), which had good agreement between the design model in the reshaped region and either the lowest energy Rosetta ab initio decoy structure or the top-ranking (by pLDDT) AF2 prediction (Cα RMSD < 2Å). All designs selected had high RMSD to the original *holo* state in the reshaped region (Cα RMSD > 4.5Å).

Yeast display screen for stable single state designs

Gene fragments of selected designs flanked by homology regions (5’: ggagggtcggcttcgcatatg, 3’: ctcgagggtggaggttccgaacaacagcttatttctgaagaggacttgta) were ordered from Integrated DNA Technologies. 10ng of synthetic DNA was amplified in a 25μLreaction using Q5 Hot Start High-Fidelity DNA Polymerase (NEB #M0494) for 30 cycles by PCR. A strong single band at the expected size was observed for all fragments. The reaction product was cleaned using Zymo Research DNA Clean & Concentrator kits for transformation of EBY100 yeast using the protocol described in (*33*). We used a modified version of the pETcon vector known as pETcon v3 (RRID: Addgene_41522) that had been altered to (i) remove a long single-nucleotide stretch near the cloning region, (ii) add an N-terminal 6x-His tag to normalize intact protein level to surface display level, and (iii) include constitutively fluorescent protein Venus as the dropout sequence to aid in colony selection (GenBank accession number: PQ010744). Single colonies were picked, inoculated into 5mL of SD-CAA medium, and incubated at 30°C overnight for each design. The starter culture was diluted 1:10 into SG-CAA to induce protein production. A volume corresponding to 5x10^5^ cells was added to a 96 well plate and spun at 3000g for 20min to pellet the cells. Cells were washed and resuspended in PBS then incubated with anti-c-Myc Mouse mAb (Alexa Fluor 647 conjugate) (Cell Signaling Technology #2233) and 6x-His Tag Antibody Alexa Fluor 488 conjugate (Fisher Scientific #MA1-135-A488) shaking at room temperature for 30min. Excess unbound antibody was washed away with chilled PBS + 1% BSA and analyzed using a Beckman Coulter Cytoflex flow cytometer. Events were gated by forward scattering area and back scattering area to collect the live cell population then by forward scattering width and forward scattering height to select individual cells for further analysis. Cells were then gated by fluorescence intensity where the threshold separating displaying from non-displayed cells was set such that 1% of an uninduced, unstained control would pass the gate. The fraction of anti-c-Myc positive cells from single cell events was used to estimate the level of expressed intact protein level on the surface of yeast as a proxy for thermostability (*34*).

Protein expression and purification for experimental screening

Plasmids (pET-28a(+)) encoding the selected designed proteins were ordered from Twist Bioscience. The DNA sequences of the designed proteins were inserted between the NdeI and XhoI restriction sites. This construct included an N-terminal 6x-His tag and thrombin cleavage site (MGSSHHHHHHGLVPRGSHM). Escherichia coli BL21(DE3) cells were transformed with these plasmids. Colonies were inoculated into 5mL LB medium and cultured at 37°C overnight. Starter cultures were diluted 1:100 into 1L of fresh LB medium and cultured at 37°C until the OD600 reached 0.6-0.8. Then IPTG was added to a final concentration of 300μM to induce protein expression at 37°C overnight. Cell cultures were centrifuged at 8000g for 10min to spin down the cells. Cell pellets were then lysed by resuspending in 4mL/g pellet of B-PER (Thermofisher #78243) with a dissolved cOmplete protease inhibitor cocktail EDTA-free tablet (Roche #COEDTAF-RO) and incubating at room temperature for 15min. Cell lysate was centrifuged at 18000g for 30min to separate the soluble and insoluble fractions. The soluble fraction was mixed with an equal volume of equilibration buffer (50mM Tris pH 7.5, 300mM NaCl, 10mM imidazole) and 1mL HisPur Ni-NTA resin slurry volume (Thermo Scientific #88222) to pull down the His-tagged proteins. Ni-NTA resin beads were washed 3 times with wash buffer (50mM Tris pH 7.5, 300mM NaCl, 25mM imidazole). For screening assays (e.g. size exclusion chromatography, dynamic light scattering, circular dichroism spectroscopy), the protein was then eluted 3 times with 0.5mL elution buffer (50mM Tris pH 7.5, 300mM NaCl, 250mM imidazole). Purity and amount of designed protein was estimated by Coomassie-stained SDS-PAGE throughout purification (BIO-RAD #456-1095).

Analytical size exclusion chromatography

Protein samples purified by His-tag pull down were analyzed using a Superdex 75 10/300 GL (Cytiva #29148721) (switch designs I89 and S89) or the Superdex 200 10/300 GL (Cytiva #28990944) (single-state designs #6306 and wildtype 1SMG) size exclusion column with 100mM KCl (pH 6.7) (Fig. S15, Fig. S16).

Dynamic light scattering

Size measurements of protein samples purified by His-tag pull down using dynamic light scattering were measured on a Zetasizer Nano S90 (Fig. S16). Samples were filtered (0.22μm) prior to measurement to remove large particles. Number distribution of hydrodynamic diameter was plotted to provide an estimate of particle size.

Circular dichroism spectroscopy

Circular dichroism (CD) data were collected on a Jasco J-710 spectrometer. Purified designs were diluted to a final salt concentration of 10mM KCl (pH 6.7). The concentrations of diluted samples were approximately 2μM determined by a Bradford assay. CD spectra were measured using a 1mm cuvette at 25°C. Melting curves were recorded at 208nm from 25°C to 95°C using a rate of 1°C/min (Fig. S15).

Protein expression and purification for NMR structure calculation

^15^N and ^13^C labeled proteins were expressed by inoculating E. coli colonies into 5mL M9 minimal medium that included 4g/L U-^13^C-glucose (99%) and 0.5g/L U-^15^NH_4_Cl (99%), which were grown at 37°C overnight. Starter cultures were then diluted 1:50 into 500mL fresh ^15^N ^13^C labeled M9 minimal medium and grown at 37°C until the OD600 reached 0.6-0.8. Then IPTG was added to a final concentration of 300μM to induce protein expression at 37°C overnight. The expressed proteins were purified by following the Ni-NTA resin pull down protocol described in the protein purification section with the following modifications: Rather than eluting the His-tagged protein from the beads, the beads were washed with and resuspended in thrombin cleavage buffer (20mM Tris-HCl, 150mM NaCl, pH 8.4). 4U of biotinylated thrombin (Novagen #696022) per 1mg of protein was added to the beads, and the immobilized His-tagged protein was cleaved from the beads while rocking at room temperature for 20h. To improve purity of the cleaved protein, the beads were pelleted, and the supernatant was added to fresh Ni-NTA resin. The beads were pelleted once more, and the supernatant was mixed with 32μL of streptavidin agarose slurry (Novagen #696022) per U of thrombin for 30min at room temperature. The sample was then filtered (0.45μm) and incubated with 100μM EDTA for 5min at room temperature to remove residual amounts of Ca^2+^. The sample was then buffer exchanged into 100mM KCl pH 6.7 by dialysis overnight at 4°C and concentrated to approximately 180μL for NMR experiments. Ca^2+^ was then added (if applicable) to the desired concentration.

Structure determination by NMR

5% D_2_O was added to samples. The final protein concentrations determined by Bradford assay were approximately 400μM. NMR spectra were all measured at 298.1K. Two dimensional (2D) ^1^H,^15^N-HSQC (pulse program: fhsqcf3gpph), 2D ^1^H,^13^C-HSQC (pulse program: hsqcetgpsisp2), 36ms 3D HCCH-TOCSY (pulse program hcchdigp3d) and 120ms 3D simultaneous ^13^C/^15^N-NOESY-HSQC (pulse program: noesyhsqcgpsismsp3d) spectra were measured using a Bruker Avance I 800 MHz spectrometer with a 5mm TCI H&F-C/N-D CryoProbe. 3D CACB(CO)NH (pulse program: hncocacbgpwg3d), 3D CACBNH (pulse program: hncacbgpwg3d), 3D Hcc(co)NH (pulse program: hccconhgpwg3d2), 3D Cc(co)NH (pulse program: hccconhgpwg3d3), 2D HBCBCGCDHDGP (pulse program: hbcbcgcdhdgp), and 2D HBCBCGCDCEHEGP (pulse program: hbcbcgcdcehegp) were collected on a Bruker Avance 600 MHz spectrometer with an Inverse 5mm H-C/N-D cryoprobe. Spectra were processed in TopSpin 3.6.3. For the Ca^2+^-bound structure, we also collected the following experiments on the Bruker Avance 600 MHz spectrometer: 2D TROSY (trosyargpphwg) and 3D ^1^H-^13^C NOESY-TROSY (noesytrosyargpphwg) to aid in the assignment of aromatic nOes. Automated peak picking, resonance assignment, and structure calculations were performed by ARTINA (*35*), and dihedral angle restraints were generated by TALOS (*36*). Manual inspection of ARTINA chemical shift assignments was done in CCPNMR v. 3.1.0 and corrections were made as necessary (*37*). The manually curated chemical shift list was then included as input for a subsequent structure calculation run in ARTINA. The output candidate structure (an ensemble of n=20 conformers) with the lowest CYANA target function value was refined in XPLOR-NIH-3.7 with the refine.py script included in the distribution (eginput/gb1_rdc/refine.py) (*38*). The 20 lowest scoring structures out of 100 were then refined in explicit water with the wrefine.py script (eginput/gb1_rdc/wrefine.py). The ensemble of the refined structures was validated using the PDB validation server (*39*). For the apo I89 structures (i.e. proposals #1 and #2 from ARTINA), we re-calculated the structures in CYANA after excluding a small subset of distance restraints that could be unambiguously assigned to the other state prior to refinement. For the Ca^2+^ bound I89 structure, the structure was first calculated with no Ca^2+^ restraints using ARTINA as described above. The Ca^2+^ restraints were then included during refinement in XPLOR-NIH-3.7, which were set to 2.8Å between the Ca^2+^ ion and the oxygens involved in coordinating Ca^2+^ (*40*).

Two-state sequence design

Initially, we attempted to use Rosetta-based methods for multi-state design. We first tried a restrained convergence algorithm, which allowed each state to explore sequence space independently and encouraged convergence to a single sequence by incrementally increasing an energy bonus if the same amino acid was sampled at corresponding positions in both states (*41*). This method did not appear to work well for states varying in conformation, as it was unlikely for positions that differed greatly in solvent accessible surface area or secondary structure to converge on the same amino acid, even with an energy bonus. We also attempted to use a genetic algorithm that allowed users to input a custom fitness function as input (*42*). However, this method performed Monte Carlo optimization of the Rosetta energy function over discrete rotamer space and was consequently computationally expensive compared to rotamer-free design methods. Due to what appeared to be low *in silico* success rates, we decided to use ProteinMPNN, a deep learning-based design method capable of quickly generating tens of thousands of multi-state designs (*17*). This method has since been shown to successfully generate proteins where domains can be hinged open and closed upon binding a peptide effector (*15*).

In general, a PDB file containing both the single-state alternative state 2 conformation and the natural Ca^2+^ binding protein state 1 conformation separated by a distance much larger than the approximate diameter of either state was generated by PyMOL. State 1 was conformer 1 of the PDB ID: 1SMG NMR ensemble relaxed using Rosetta with restraints on heavy atom positions. Ca^2+^ was removed from the structure, as this implementation of ProteinMPNN did not support ligand atoms.

The state 2 conformation was generated with the following design task in mind: beyond individual bistable designs, we wanted to design a family of sequences that had significantly different state population distributions in the absence of ligand despite having high sequence identity. We reasoned it would be more likely to design such a set of sequences if we first restricted the allowable sequence space during design. Initially, the set of potential designable residues for two-state design included the reshaped region and its neighbors in both states as defined for single-state design (n=37 residues). To exclude positions that did not change in environment significantly between states or were otherwise not critical in determining the preferred structure, we individually mutated each designable position in the state 2 design to the corresponding state 1 amino acid and predicted the structural impact of each “reversion” using AF2 (though one could also use an in silico deep mutational scanning approach to evaluate other potential amino acids at each position). If the predicted structure was still consistent with the state 2 conformation (Cα RMSD < 1.5Å), this mutation was considered to be in the “tolerated” sequence space of state 1. We then input a state 2 sequence containing all “tolerated” amino acid reversions into AlphaFold2 and used the best-ranked structure by pLDDT as the state 2 state during two-state design. Of note, this prediction was still highly similar to the original single-state design in structure (Cα RMSD = 1.15Å in the reshaped helix (residues 59-69) and 1.21Å over the entire backbone) and had an average pLDDT of 84.6 in the reshaped helix. The final set of designable residues included: (1) all positions that were not reverted to the corresponding state 1 amino acid through this process and (2) their neighbors (defined as in single-state design) (n=25 residues) (Table S2).

All corresponding positions between states were tied together across chains during design with equal weighting of each state. During preliminary rounds of design, we observed the last turn of the reshaped helix (residues 68-72) fraying at the C-terminal region in AF2 structure predictions. To stabilize the helix, position 62 was biased toward residues that could hydrogen bond with the side chain of D66 in subsequent rounds of two-state design, i.e. K, R, and Q. ProteinMPNN was used to generate 10^4^ sequences, which were input without MSA generation into ColabFold for structure prediction (*43*).

NMR experiments to characterize dynamics

U-^15^N-labeled protein was prepared as described above for structure calculation except unlabeled glucose was used in place of U-^13^C-glucose (99%) in culture media. All experiments were done on the Bruker Avance I 800 MHz spectrometer. The concentration of samples ranged from 100-200μM in 100mM KCl, pH 6.7, 5% D_2_O. An initial screen for Ca^2+^ modulation of conformational dynamics was done by collecting ^1^H,^15^N-HSQC spectra of all two-state designs with and without 10eq Ca^2+^ (pulse program: fhsqcf3gpph). For design I89, a titration with 0eq, 1eq, 5eq, and 10eq of Ca^2+^ was done to track chemical shift changes compared to the assigned apo ^1^H,^15^N-HSQC spectrum. To further probe which residues may be dynamic even without the addition of Ca^2+^, a ^1^H,^15^N-HSQC temperature series from 5°C to 35°C in 5°C increments was collected for design I89 and S89. To characterize motions occurring on a low microsecond timescale, we acquired ^15^N rotating frame (R_1ρ_) relaxation dispersion experiments at T=298.1K (pulse program: hsqctretf3gpsitc3d) with a spin-lock field strength of 3.0kHz and delay times of 2, 4, 6, 8, 10, 20, 30, 50, 70, 100ms, where the 30ms experiment was repeated to assess reproducibility and estimate error. T_1ρ_ values were calculated using the Bruker Dynamics Center 2.8.4.

Molecular dynamics simulations

Simulations of designs in the presence and absence of Ca^2+^ were performed using GROMACS 2022.5 (*44*). The initial structures were the AMBER-relaxed AF2 predictions for the design with the N-terminal thrombin cleavage site scar included to be consistent with experimentally characterized designs. The proteins were parameterized using the a99SB-disp force field and water molecules were parameterized using the a99SB-disp water model (*45*). The systems were solvated and neutralized as in Table S6. Energy minimization of the system was performed with the steepest descent minimization algorithm to a tolerance of 1000.0 kJ mol^-1^ nm^-1^. Equilibration was performed in the NVT ensemble for 1000ps at 300K using the Berendsen thermostat. Systems were then equilibrated in the NPT ensemble for 100ps at a target pressure of 1 bar at 300K maintained by the Berendsen thermostat with position restraints on all heavy atoms. Bond lengths and angles of protein atoms were constrained with the LINCS algorithm and water constraints were applied using the SETTLE algorithm. The PME algorithm was used for electrostatics with a grid spacing of 1.8nm. Van der Waals forces were calculated with the Verlet cut-off scheme using a 1.2nm cut-off distance. Snapshots were saved every 80ps. Initial structures were AF2 structure predictions of two-state designs (including the residues GSHM at the N-terminus corresponding to a scar from the thrombin cleavage site) relaxed in AMBER. The Cα RMSD of the reshaped region (residues 53-76) were calculated over the course of a 1μs simulation using GROMACS 2022.5 command line tools. To quantify correlations between residue dihedral angles during the apo I89 simulation, the trajectory from 100ns-800ns was split into seven 100ns blocks. For each block, dihedral angle data was extracted with:

gmx chi -f md.xtc -s md_1_us.tpr -phi -psi -omega -rama -all -maxchi 4 -HChi -b 5000

We took six out of five segments (not necessarily contiguous) as a sample ensemble from which mutual information was calculated using the MutInf method (*23*):

dihedral_mutent -x <base dir> -d / -o 6 -n 7 -w 30 -p 0 -c “yes” -a “yes” I89.reslist > I89_mutinf.out

where the *.reslist file was a list of all residues. The file containing mutual information between pairs of residues, with zero diagonal (*bootstrap_avg_mutinf_res_sum_0diag.txt) was visualized a heatmap.

Frame2seq scoring of point mutants

To predict the effects of single point mutations on the conformational equilibrium of switch design I89, we computed sequence likelihoods given structure using Frame2seq (*18*). Frame2seq models learn to approximate P(sequence|structure) via a masked language modeling objective. Pseudo log-likelihood (PLL) has been explored for scoring sequences (*46*), which we adapted for a structure-conditioned ranking task. For each conformational state (where the AF2 models of I89 and S89 were used to represent states 1 and 2, respectively), we separately output Frame2seq model PLL by providing the structure and the sequence as input with a mask introduced at the mutated position. We then subtracted the single point mutant negative PLL from the wildtype negative PLL (the reference point) to compute the score as follows:

$$score= log(p\left( x_{i}=x_{i}^{wt} | x_{-i}^{wt},Y \right)-log(p(x_{i}=x_{i}^{mt}|x_{-i}^{mt},Y)$$

where $\boldsymbol{i}$ is the mutated position, $\boldsymbol{x}^{\boldsymbol{wt}}$ is the wildtype sequence, $\boldsymbol{x}^{\boldsymbol{mt}}$ is the mutant sequence, $\boldsymbol{x}_{\boldsymbol{-i}}$ is the sequence $\boldsymbol{x}$ with a mask introduced at position $\boldsymbol{i}$, and $\boldsymbol{Y}$ is the structure.

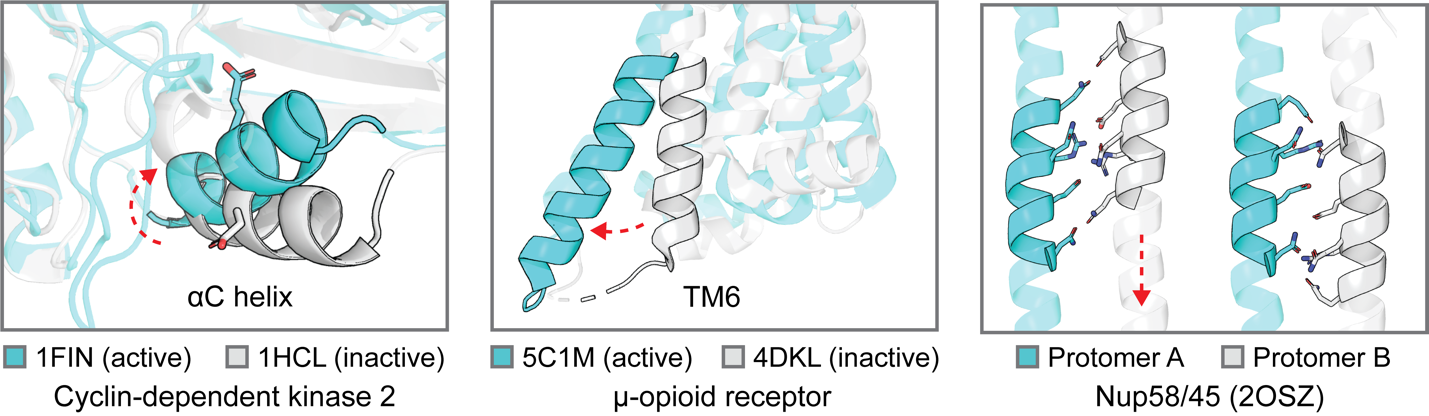

**Fig. S1: Intra-domain conformational changes are common in natural proteins and essential to control function.** For instance, helices are known to undergo several modes of motion including twisting, kinking, and sliding to perform functions like forming active catalytic sites (e.g. rotation of the αC helix in kinases, left panel), bind to downstream signaling partners (e.g. kinking of TM6 to allow for G-protein coupling in GPCRs, middle panel), and modulate pore diameters (e.g. sliding of helices in nucleoporins, right panel). PDB codes for the different conformations (grey, cyan) are indicated.

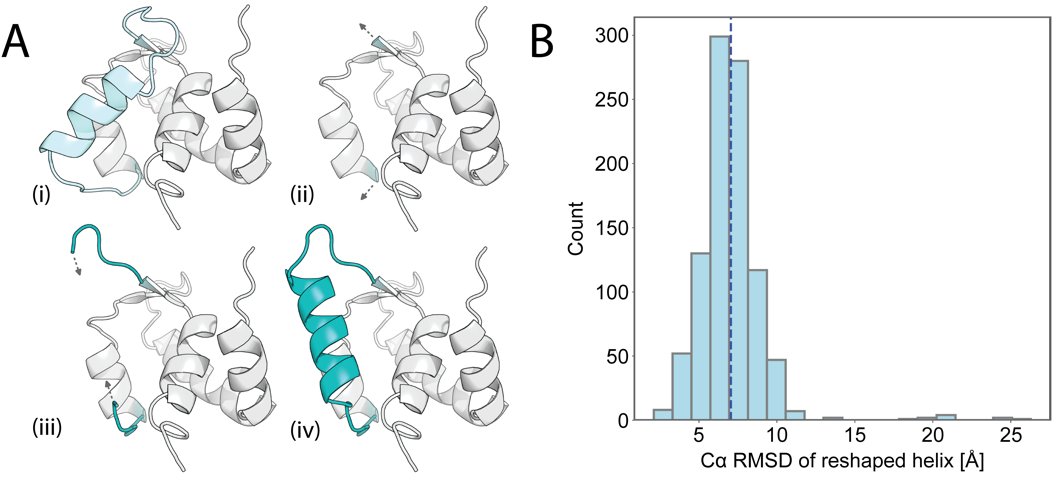

Fig. S2: Generating de novo alternative conformations. (A) The loop-helix-loop unit combinatorial sampling algorithm applied to PDB ID: 1SMG involved (i) designating a reshaped region (light cyan), (ii) removing the existing loop-helix-loop subunit, (iii) sampling new loops from the PDB, and (iv) growing helices from both loops and closing the helix (if possible) to generate a de novo loop-helix-loop conformation (teal). (B) The distribution of Cα RMSD between the starting helix conformation and the de novo reshaped helix averaged around 7.1Å (dashed line).

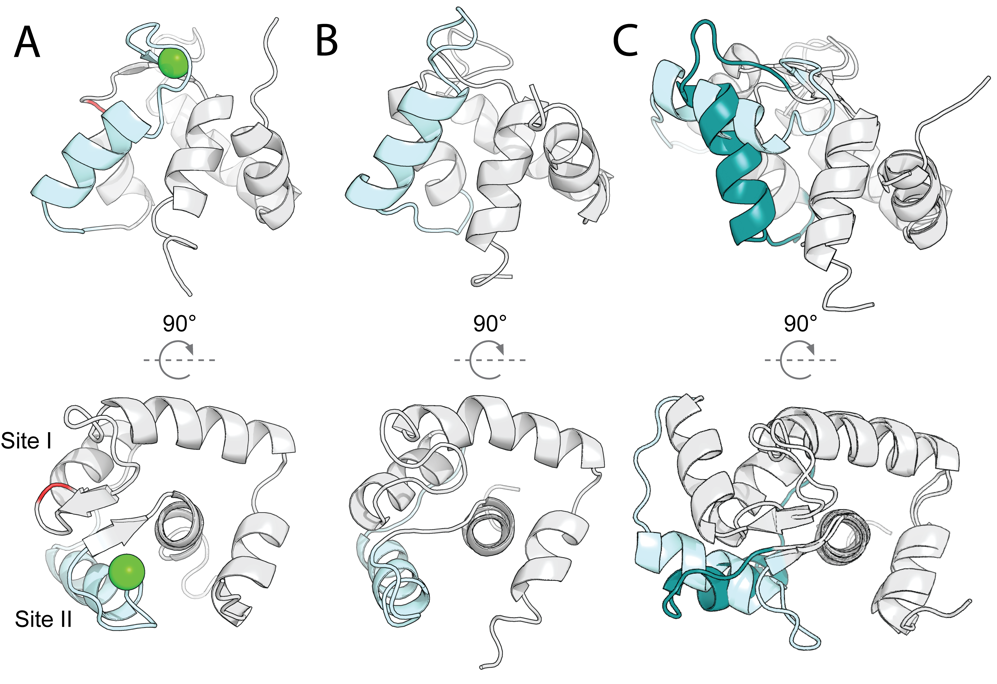

Fig. S3. Engineered Ca^2+^ binding *holo* state (state 1). (A) Structure of the E41A point mutant (red) of the N-terminal domain of skeletal muscle Troponin C from *Gallus gallus* (PDB ID: 1SMG, used as binding-competent state 1) in two orientations. While the wildtype protein has two Ca^2+^ binding loops (site I and site II) with binding affinities in the low μM range and a closed-to-open transition upon binding, the E41A mutation lowers the affinity of site I to the low mM range and decouples Ca^2+^ binding from conformational change. The conformation upon binding is now highly similar to the “closed” *apo* conformation of the wildtype protein (PDB ID: 1SKT) shown in (B) (Cα RMSD = 1.6Å, excluding loops). (C) Structure of the “open” holo conformation of the wildtype protein (pale cyan, PDB ID: 1TNX) is distinct from the de novo generated state 2 conformation designed in this study (teal, Fig. 1C in the main text, bottom row) (Cα RMSD = 6.7Å, computed over the reshaped helix).

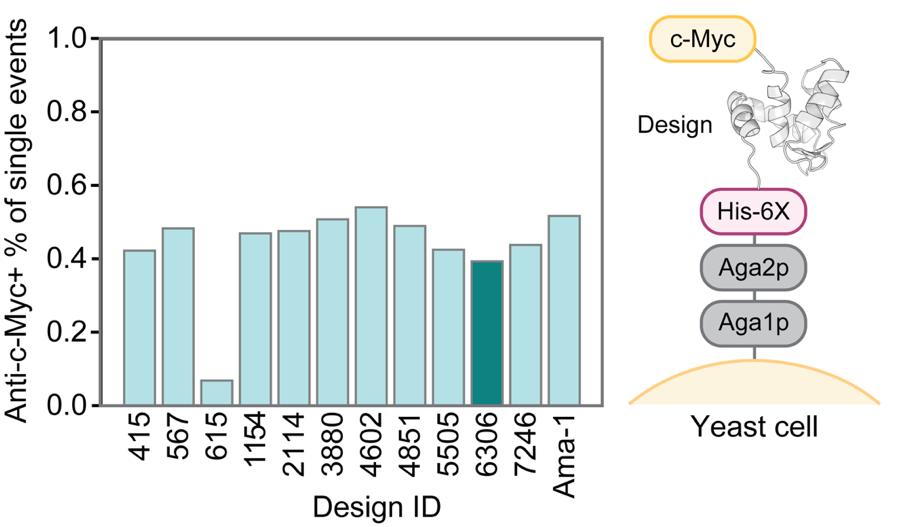

**Fig. S4: Yeast display to screen alternative (state 2) single-state designs.** Flow cytometry was used to sort cells displaying many copies of a designed protein labeled at the C-terminal c-Myc tag with a fluorescent antibody. 11 designs (design ID, x axis) were screened by surface display level of non-degraded protein (i.e. the anti-c-Myc positive percentage of single cell events), which is known to be correlated with thermostability. A highly stabilized version of the Ama-1 protein (gifted by Eric Klavins) was included for comparison. For further characterization, we selected design #6306 (teal).

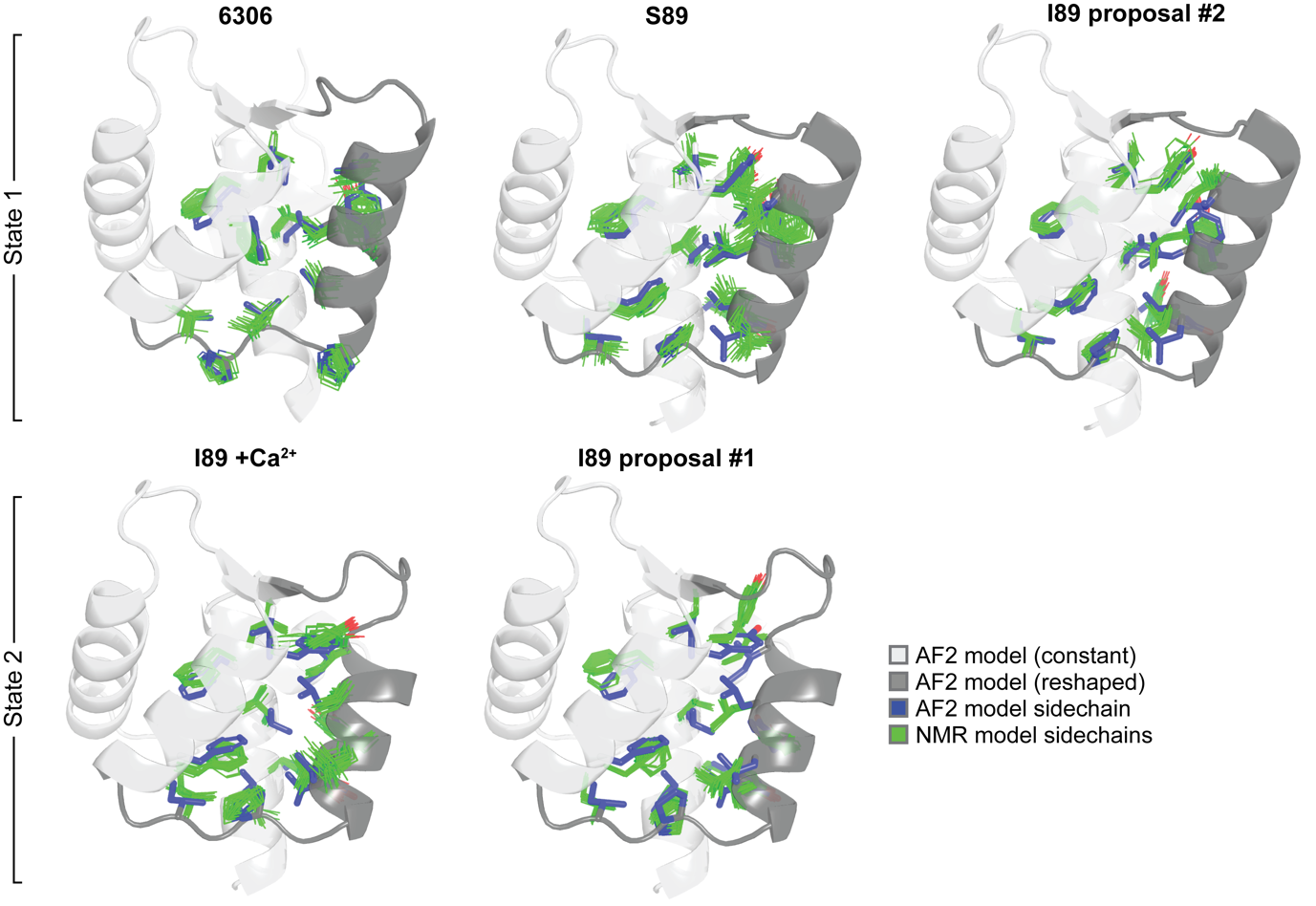

**Fig. S5: Sidechain details of hydrophobic residues.** The backbones of the designed models are shown are cartoons, where the constant region is light grey while the reshaped region is dark grey. The hydrophobic residues of each model are shown as blue sticks. The sidechains of the NMR models (n=20) are shown as green lines. Overall, there is good agreement, in particular for design #6306 (single-state design) and design S89. For the I89 models we observe heterogeneity due to the presence of multiple states coexisting in solution (i.e. the NMR structures are time and ensembled-averaged representations of a multi-state system, resulting in models that do not necessarily represent the individual states sampled by I89 at equilibrium). However, the backbone conformations and characteristic local environments for key residues TYR 43/ILE 69 for apo I89 NMR structure proposal #1 and proposal #2 were in line with states 1 and 2, respectively (Fig. 2D). Upon addition of Ca^2+^, I89 adopts a conformation closer to state 1 but still samples state 2 to some extent according to the assigned distance restraints (Fig. S12).

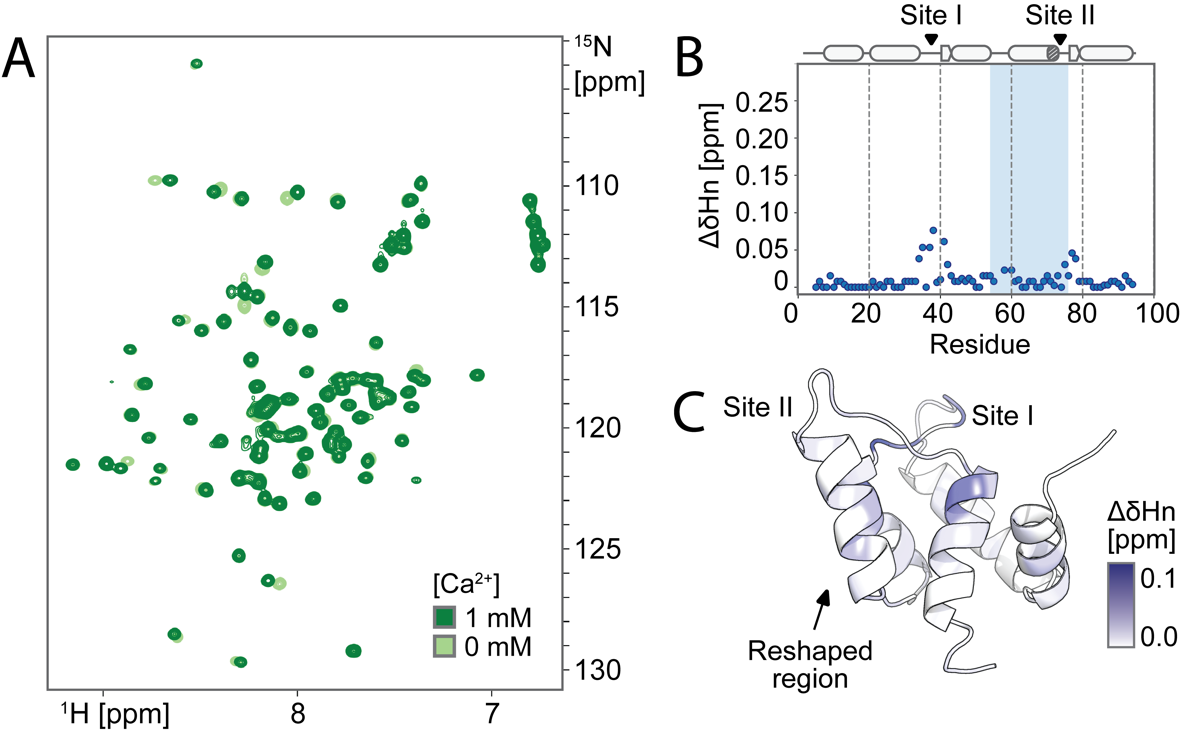

**Fig. S6:** **The single-state design for the state 2 (binding-incompetent) conformation does not undergo structural change upon binding Ca^2+^.** (**A**) ^1^H,^15^N-HSQC spectra of the single-state design #6306 (state 2) +/- 10 eq. Ca^2+^ revealed chemical shift changes localized primarily to the unchanged weak binding site I. (**B**) While there were small chemical shift changes in site II, the reshaped region (residues 53-76, shaded) remained essentially unchanged, which is also (**C**) visualized structurally, suggesting Ca^2+^ did not induce structural change to a 1SMG-like conformation in the single-state design.

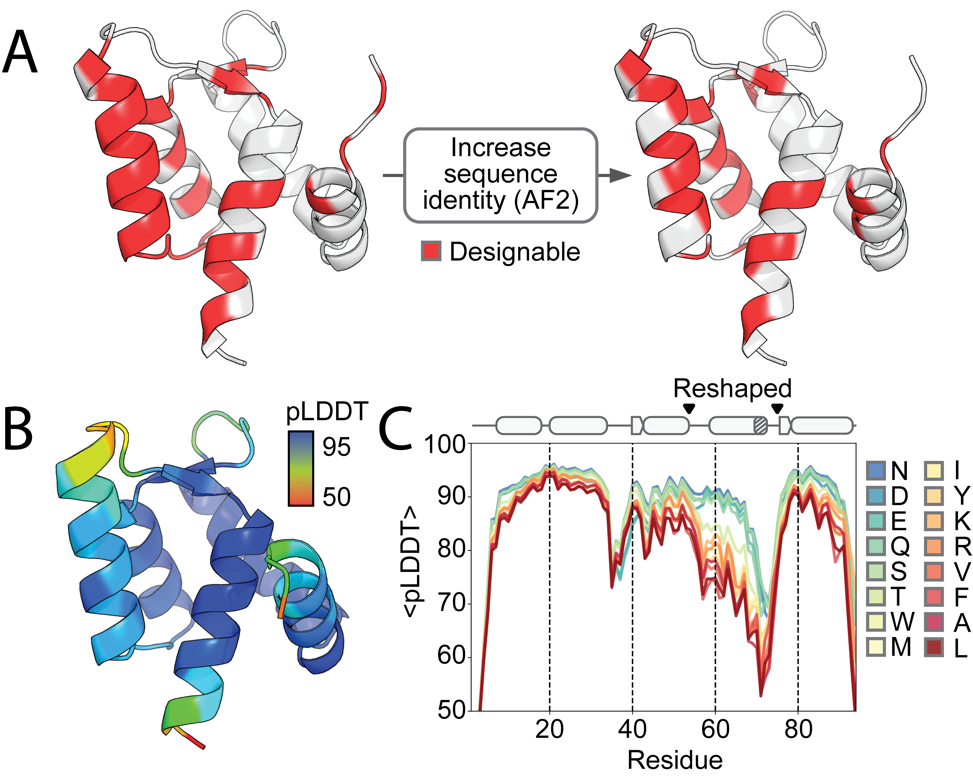

**Fig. S7: Generating the high sequence identity alternative state (state 2) for multi-state design.** (**A**) AF2 model of the original single-state state 2 design, colored by designable residues during multi-state design before (left) and after (right) sequence identity to state 1 was increased by the method detailed in Fig. 1B. The final set of designable residues only included positions that appeared to impact state preference according to AF2 (Table S1). (**B**) The AF2 model of the apo sequence with all “tolerated” mutations that increased sequence identity combined colored by pLDDT. The AF2 prediction was in still in excellent agreement with our original state 2 model (Cα RMSD = 1.15Å in the reshaped helix (residues 59-69) and 1.21Å over the entire backbone). (**C**) The average pLDDT of the five AF2 models for each point mutant at position 89.

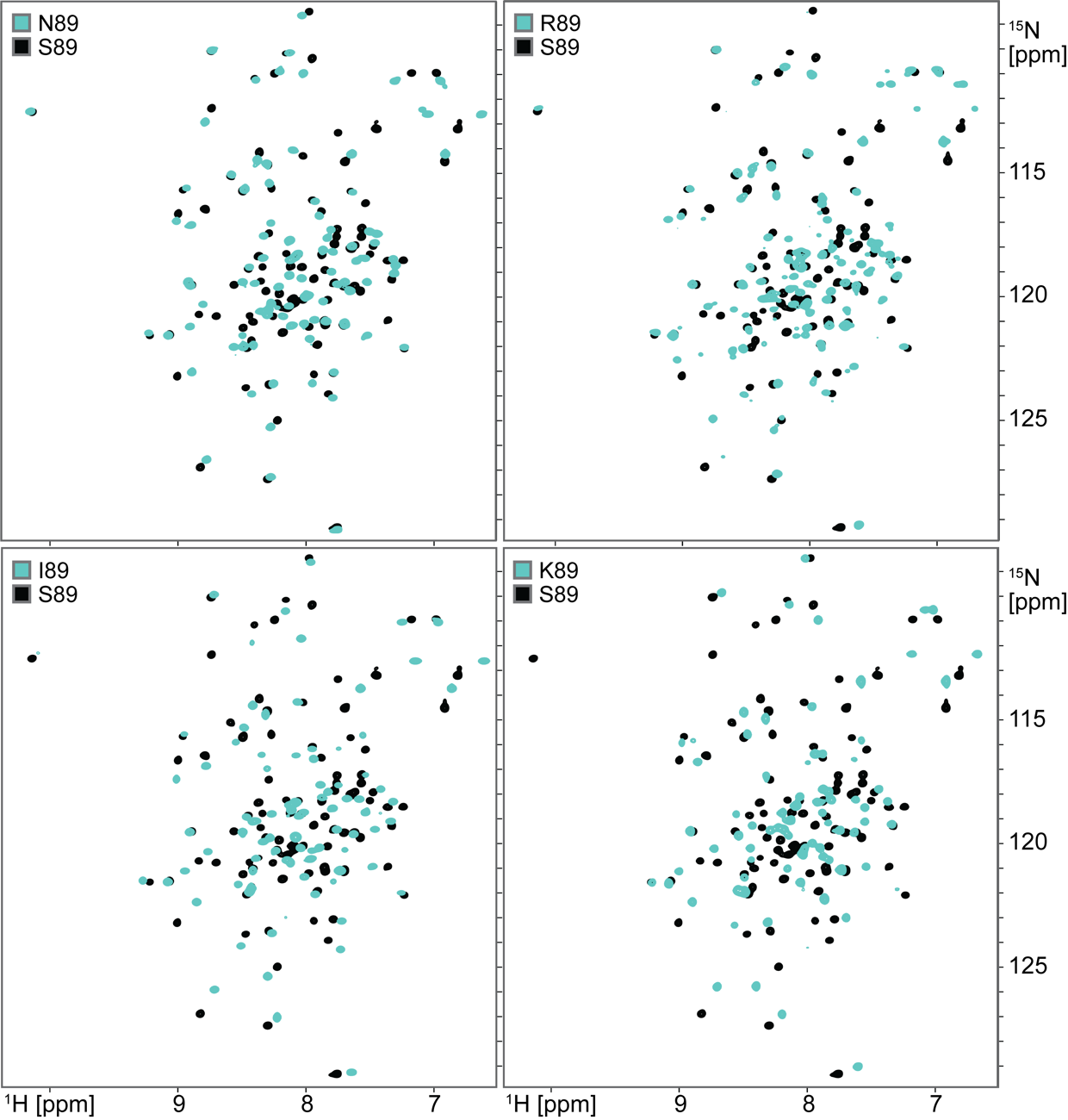

**Fig. S8 Comparison of *apo* ^1^H,^15^N-HSQC spectra of switch designs with allosteric point mutations.** The ^1^H,^15^N-HSQC spectra for designs N89, R89, I89, and K89 (teal) are shown overlaid with that of S89 (black) for reference. Despite only differing at residue position 89, all spectra show widespread chemical shift differences, indicating changes in the conformational ensembles between designs in regions outside of just the mutated site. As the spectra for designs R89 and K89 showed greater evidence of line broadening (potentially as a result of exchange on an intermediate timescale), we focused structural studies on designs I89 and S89 as extremes (by chemical shifts) of the observed two-state equilibrium. We note that the NMR spectra for all five designs show evidence of two-state exchange, while AF2 predictions showed pronounced structural variation only for R89 (Fig. 2A).

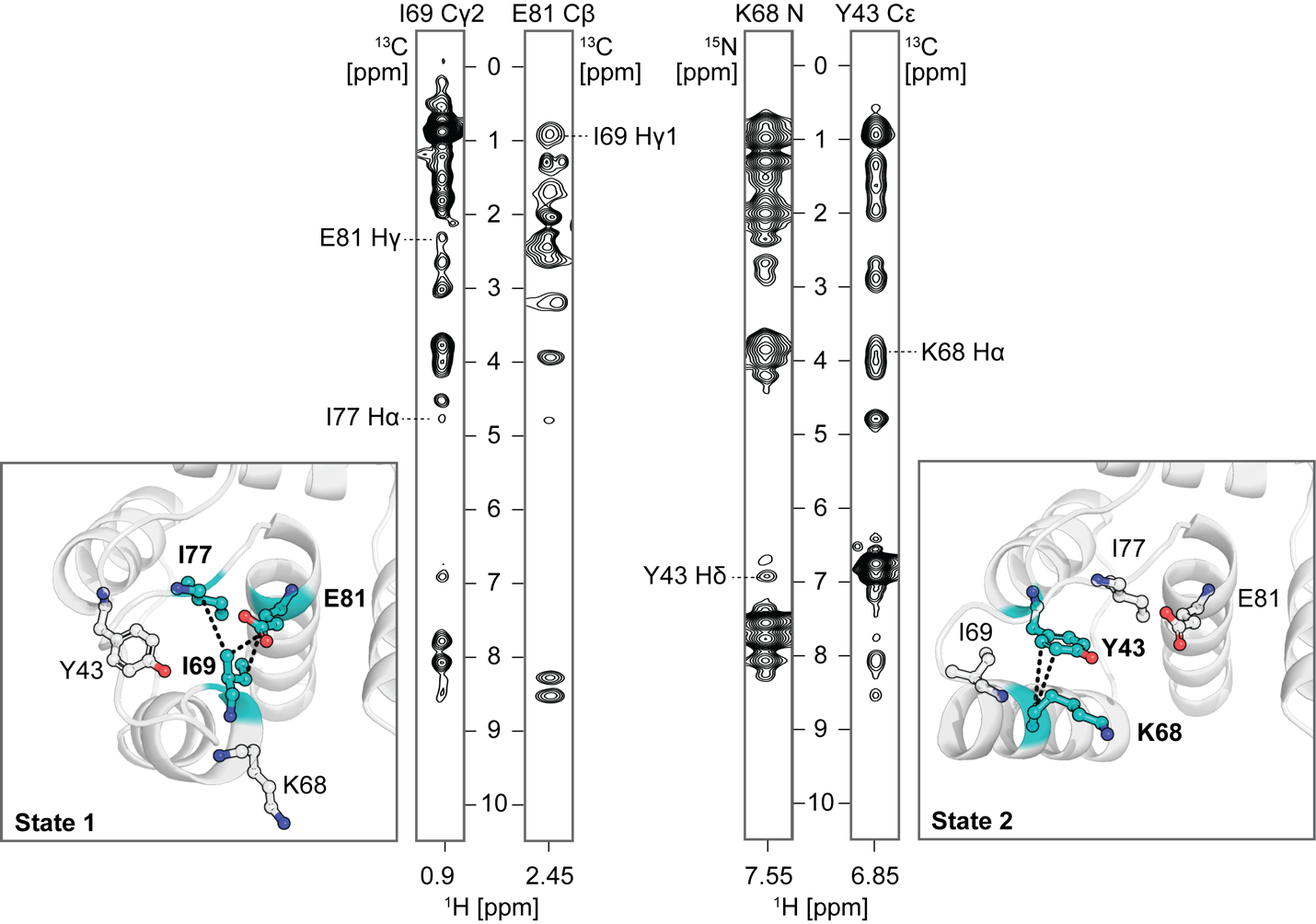

**Fig. S9: Distance restraints consistent with both states 1 and 2 were assigned for *apo* I89 by ARTINA.** For NMR structure proposals #1 and #2, ARTINA assigned distance restraints that were consistent with states 1 and 2, respectively. Representative NOESY strips containing assigned peaks unique to one state are shown. Distance restraints are also visualized structurally (panels, relevant residues in close contact for each state are bolded and colored in teal).

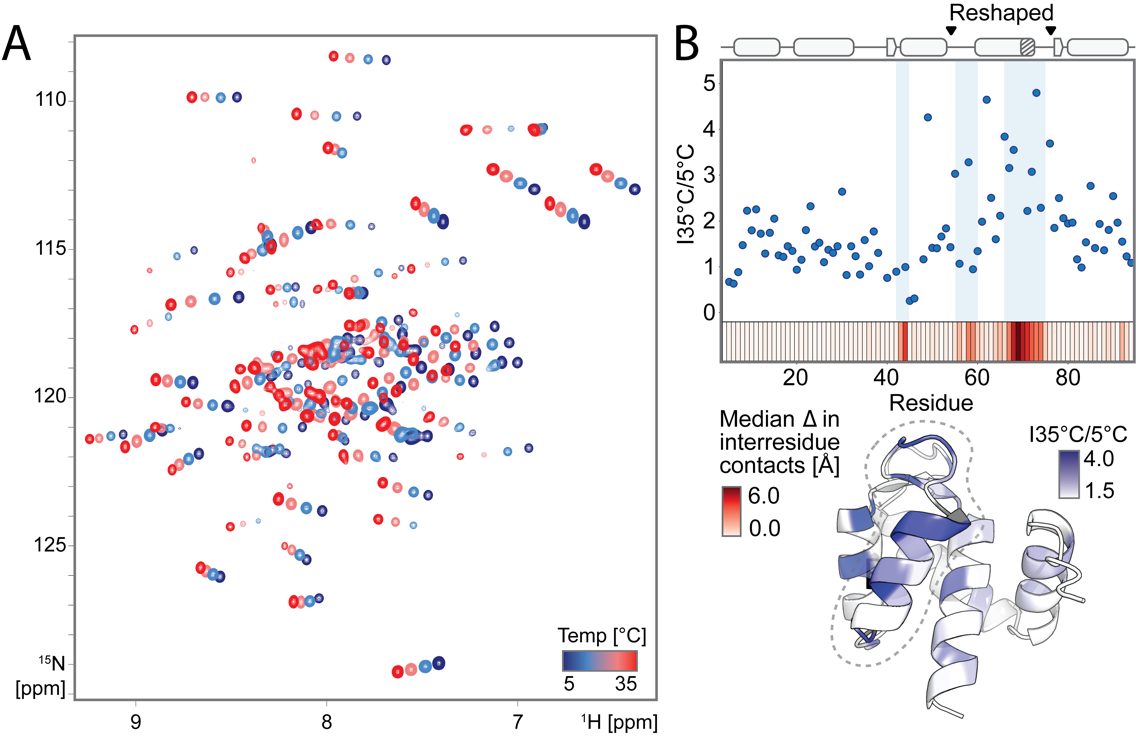

**Fig. S10: ^1^H,^15^N-HSQC temperature series for multi-state switch design I89.** (**A**) Overlaid ^1^H,^15^N-HSQC spectra for I89 taken at 5°C intervals from 5°C to 35°C. (**B**) The ratio of peak intensity at 35°C to 5°C visualized on the AF2 model (bottom) for I89 and by residue (top) shows elevated changes to intensity with temperature in regions predicted to undergo significant conformational exchange. Expected changes to the local chemical environment of residues (as measured by the median change in Cβ-Cβ distances of close contacts between states) are shown on the x axis in the top plot where darker values indicate larger expected changes. In particular we note lower intensity at lower temperatures, consistent with a transition from a fast to an intermediate exchange regime. We observe notable agreement between our temperature series data and our R_1ρ_ relaxation rate measurements (shown in Fig. 2E in the main manuscript).

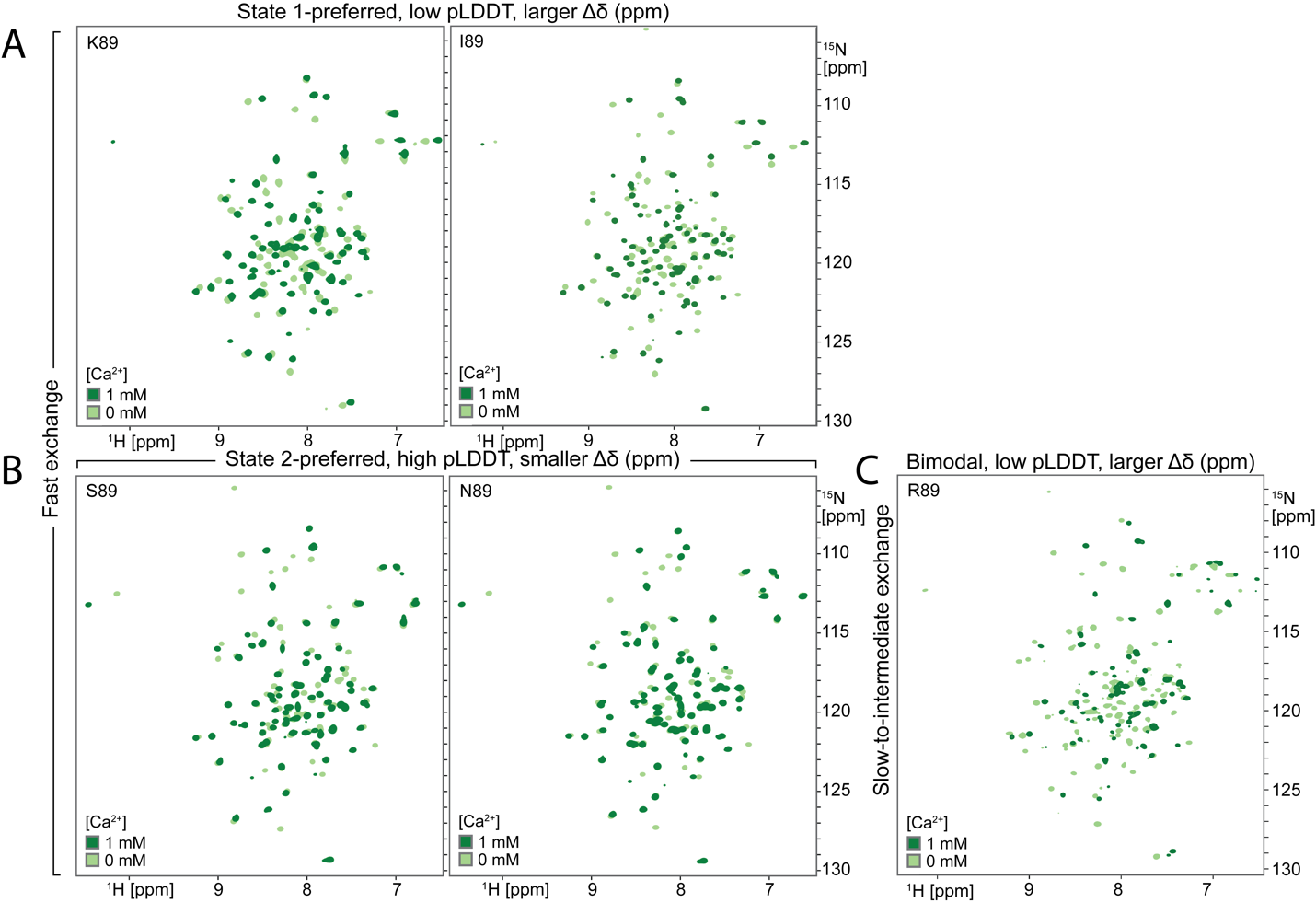

**Fig. S11: Effect of Ca^2+^ addition on ^1^H,^15^N-HSQC spectra of switch designs with allosteric point mutations.** The addition of approximately 10eq. of Ca^2+^ to point mutant designs caused changes in chemical shift to a large number of peaks in the ^1^H,^15^N-HSQC spectrum. (**A**) State 1 preferred (I89 and K89) designs, as predicted by AF2, had larger chemical shift changes in magnitude overall compared to (**B**) state 2 preferred designs (S89 and N89), which should primarily adopt a binding-incompetent conformation. (**C**) Design R89 had a considerable number of minor peaks both with and without Ca^2+^. Additionally, many peaks decreased in intensity upon Ca^2+^ addition, suggesting conformational exchange on slow and intermediate timescales, respectively.

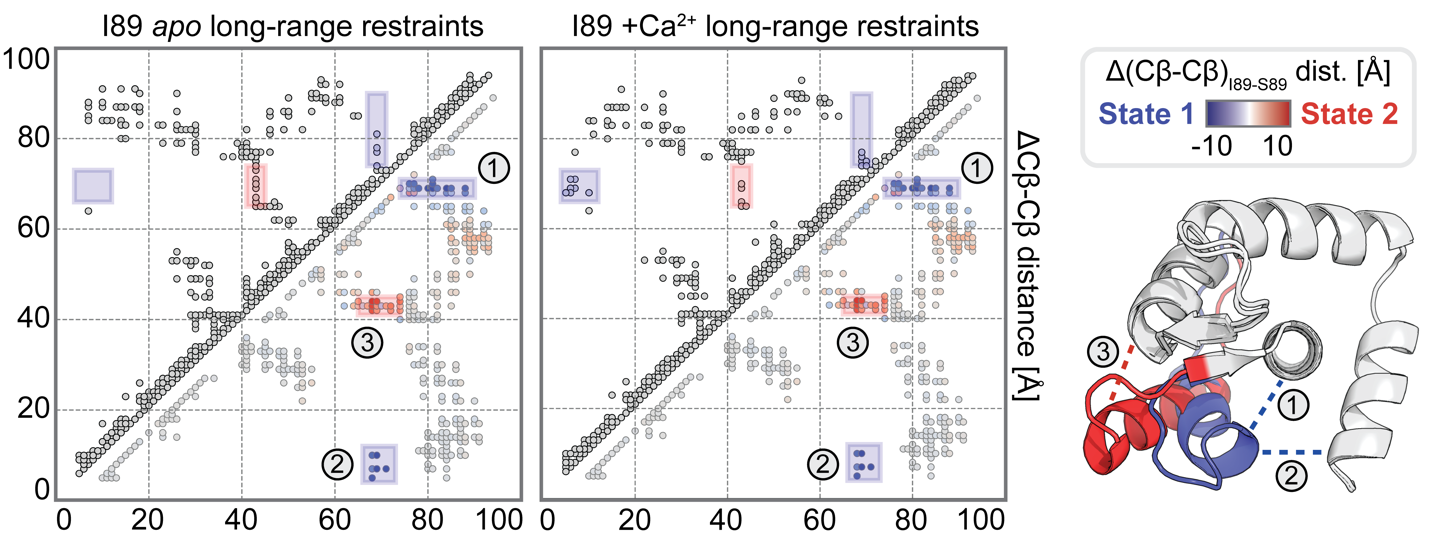

**Fig. S12: Distance restraints become more consistent with state 1 upon Ca^2+^ addition.** Comparing assigned nOe-derived distance restraints (upper triangles) with a difference contact map (lower triangles) shows a shift in state with the addition of Ca^2+^. Specifically, distance restraints characteristic of both state 1 (blue) and state 2 (red) are assigned for the apo protein. Adding 10eq. of Ca^2+^ results in many more assigned restraints unique to state 1 and fewer restraints unique to state 2, in line with Ca^2+^ preferentially stabilizing state 1.

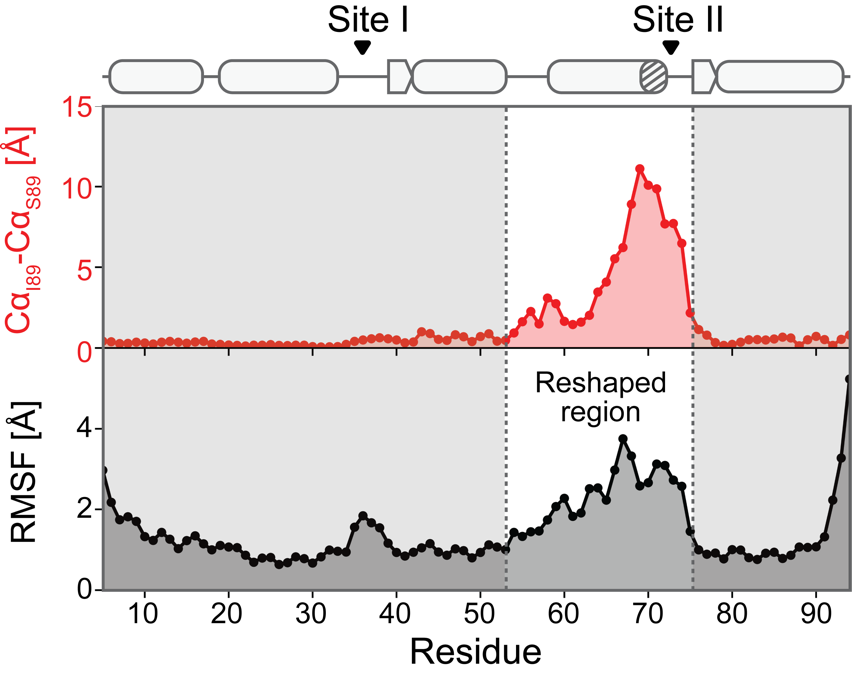

**Fig. S13: Root mean square fluctuation (RMSF) of apo I89 per residue.** The upper plot (red) shows the distance between corresponding Cα atoms of the AF2 predictions for I89 and S89 (after superimposing on the non-reshaped portion of the backbone). The bottom plot (black) shows the RMSF calculated for the apo I89 trajectory from 0.1 to 0.8μs (consistent with the timeframe used for our mutual information analysis) As expected, we saw higher fluctuations associated with an increase in expected conformational change. The other small increase in observed fluctuations near residues 35-40 is associated with the non-reshaped lower affinity Ca^2+^ binding site I.

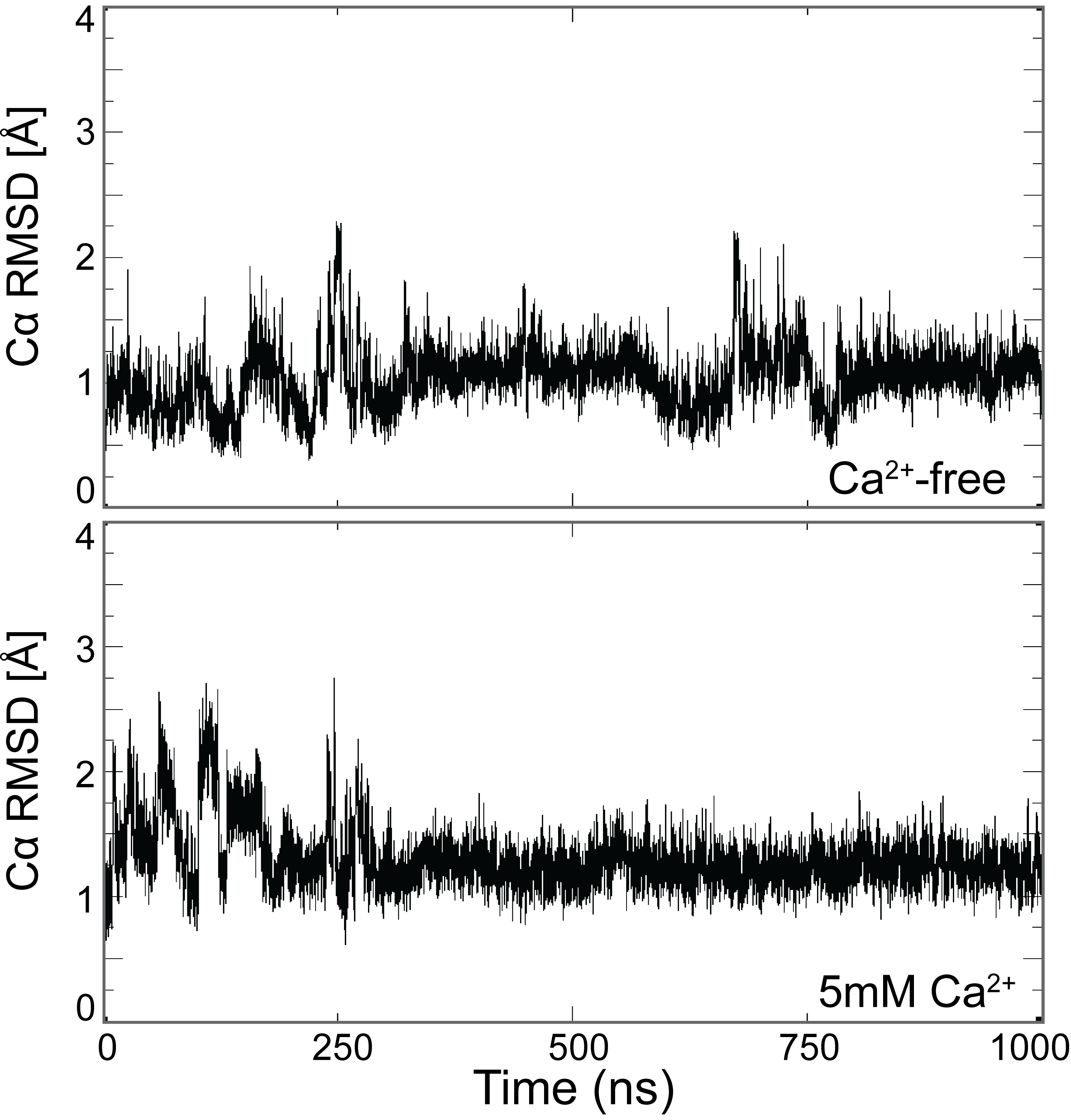

**Fig. S14: Molecular dynamics simulations for design S89.** The plots show the Cα RMSD of the reshaped helix compared to the initial conformation over the course of a 1μs simulation with (bottom) and without (top) Ca^2+^. In both simulations, the reshaped helix is in a conformation consistent with state 2 (with small fluctuations) for the entire trajectory without sampling state 1. This result is consistent with the relatively high (confident) average pLDDT of design S89 in the reshaped region.

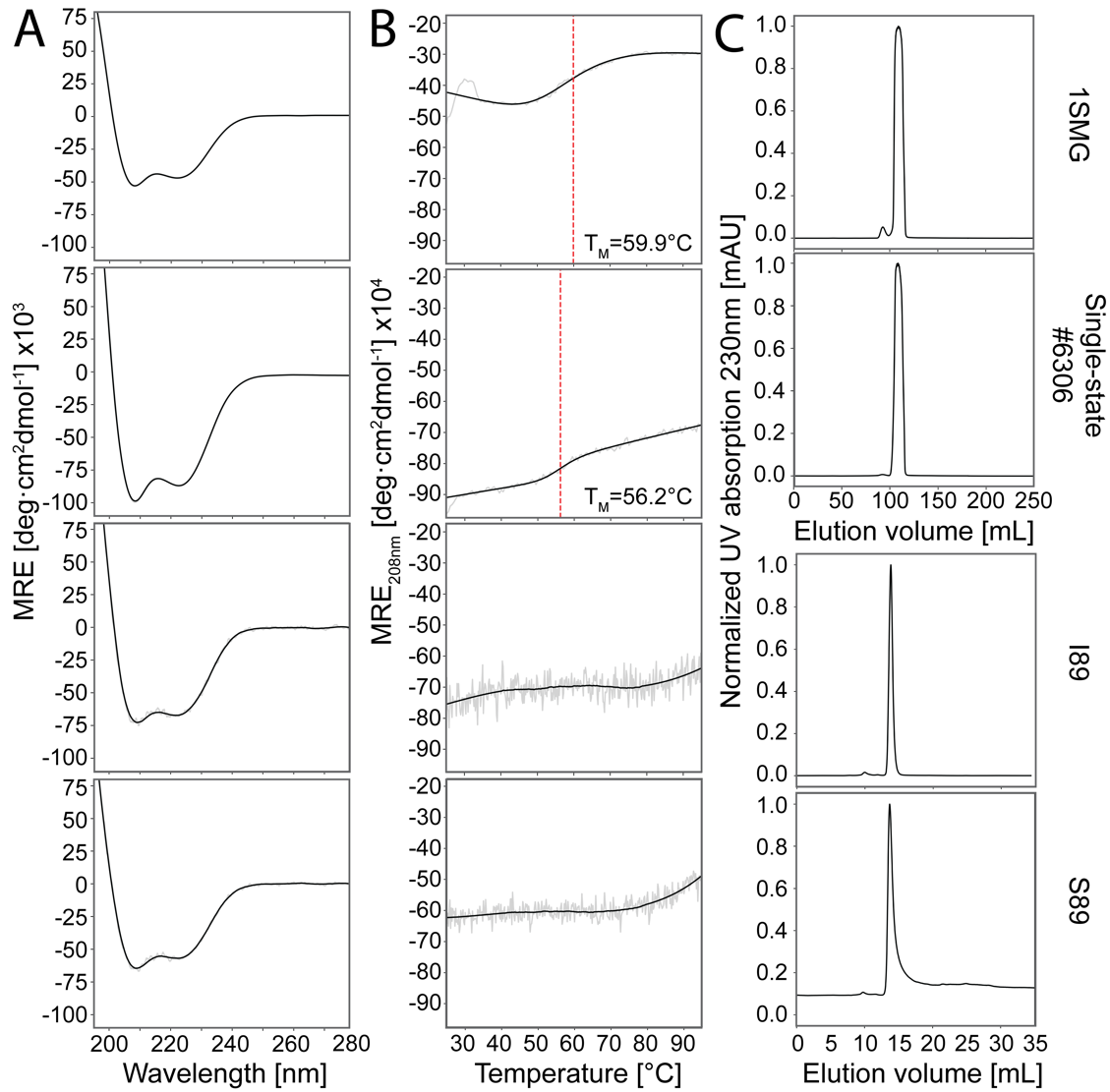

**Fig. S15: Biophysical characterization of single-state and switch designs.** (**A**) Circular dichroism spectra of 1SMG (state 1), single-state apo design #6306 (state 2), switch design I89, and switch design S89 from top to bottom, respectively, were all consistent with α-helical proteins. (**B**) Changes in the mean residue ellipticity measured at 208nm over increasing temperature reveal moderate melting temperatures for 1SMG (a natural protein) and single-state apo design #6306 (RosettaDesign), compared to the highly stable ProteinMPNN switch designs I89 and S89. (**C**) Size exclusion chromatography (SEC) curves show a predominantly monodisperse species for all designs.

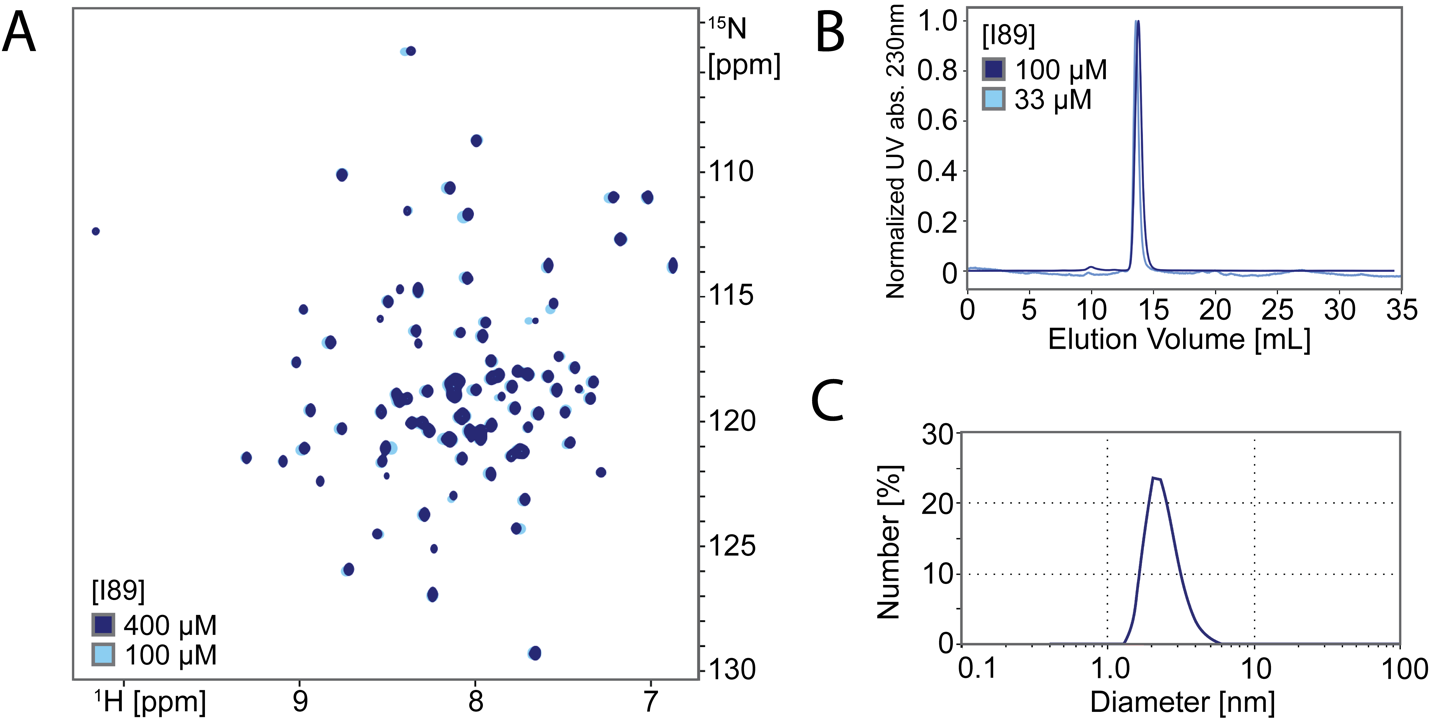

**Fig. S16: Verifying the oligomeric state of switch design I89.** (**A**) ^1^H,^15^N-HSQC spectra of I89 do not show significant changes in chemical shift or peak intensity with varying protein concentration. (**B**) The SEC curves of I89 similarly do not display shifts to smaller elution volumes (i.e. larger hydrodynamic volumes) at higher protein concentrations. (**C**) The size distribution by number measured by dynamic light scattering of I89 at 400μM shows a unimodal distribution centered around ~25Å in diameter, corresponding to the approximate size of monomeric I89. Taken together, these results suggest that I89 is predominantly in a monomeric state at the concentrations used in this study.

| Position (State 1 / State 2) | Structural environment | Notes |
| --- | --- | --- |
| 46  LEU / PHE | 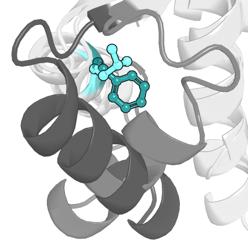 | In helix B facing the reshaped helix (helix C): important for packing of helix B and reshaped helix against each other and protein core |
| 55  GLN / VAL | 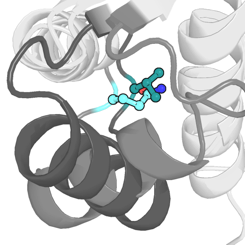 | In loop III: facing the central core helix (helix D): important for packing of helix B and the reshaped helix against the protein core |
| 58  THR / ASP | 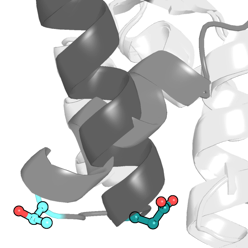 | In loop III: solvent-exposed in state 1, facing the central core helix in state 2; hydrogen-bonding from the reshaped helix to the central core helix in state 2; sterically constrained in state 2 |
| **61**  **GLU / VAL** | 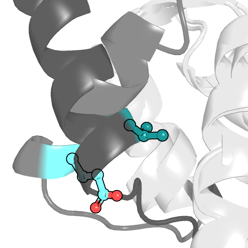 | In reshaped helix: large change in solvent-accessible surface area (SASA) (44% to 1% from state 1 to state 2); sterically constrained in state 2 |
| 62  LEU / GLN | 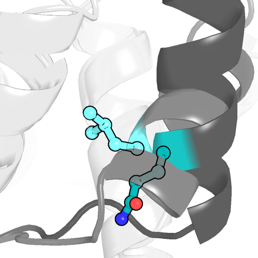 | In reshaped helix: intermediate accessibility in state 1, forms hydrogen bond with loop III in state 2; important for orienting reshaped helix in state 2 |
| 64  ALA / TYR | 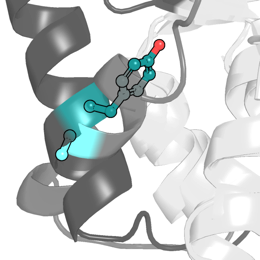 | In reshaped helix: solvent-exposed in state 1, forms hydrogen bond with central core helix and packs against protein core in state 2 |
| **66**  **ILE / LYS** | 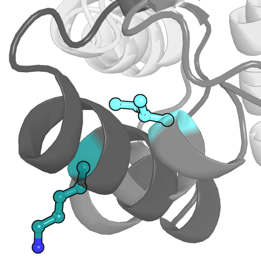 | In reshaped helix: large change in SASA (15% to 66% from state 1 to state 2) |
| **68**  **GLU / LEU** | 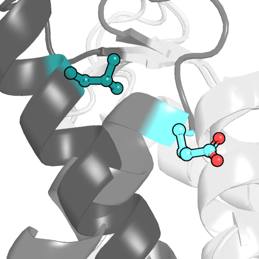 | In reshaped helix: forms hydrogen bond to central helix in state 1, packs against the hydrophobic core in state 2; important for orienting the reshaped helix in both state 1 and state 2 |
| 71  GLU / GLN | 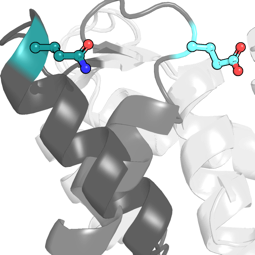 | In Ca^2+^ binding site II: hydrogen bonds with the central helix and the N-terminal helix (helix N) in state 1, solvent-exposed in state 2 but seems to impact propensity of the reshaped helix to fray at the C-terminus and form a β-hairpin |
| 85  MET / LEU | 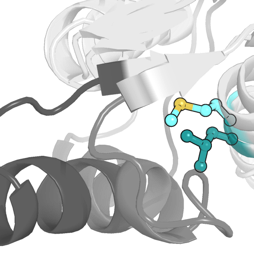 | In central core helix facing the reshaped helix: important for orienting the reshaped helix in both states |
| **89**  **GLN / SER** | 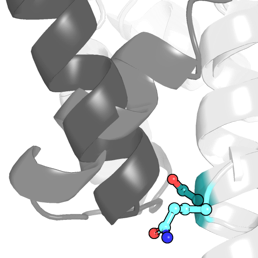 | In central core helix facing loop III: hydrogen bonds with loop III in state 2; sterically constrained in state 2 |

**Table S1: Residue positions key for determining state preference.** As described in Fig. 1B, we first evaluated mutations were mutating the amino acid in state 2 to the corresponding amino acid in state 1. The single-state design for state 2 with greater sequence identity to state 1 resulting from this process was used in multi-state design. We also performed an in silico deep mutational scan of all amino acids (except proline, glycine, histidine, and cysteine) at each position to increase sequence identity further, and the 4 remaining positions that could not be mutated to the same amino acid in states 1 and 2 without perturbing the predicted structure are bolded. State 1 is colored in light grey (reshaped region) and light cyan (amino acid), while state 2 is colored in dark grey (reshaped region) and teal (amino acid). Residues predicted to be key to determining state preference were typically positions with changes to hydrogen bond networks, sterics, or solvent accessible surface area (SASA) between states.

| **Design** | **Sequence** | **# mutations** | **Designable residues** |
| --- | --- | --- | --- |
| 1SMG  (state 1 single-state) | ASMTDQQAEARAFLSEEMIAEFKAAFDMFDADGGGDISTKALGTVMRMLGQNPTKEELDAIIEEVDEDGSGTIDFEEFLVMMVRQMKEDA | 0 | -- |
| 6306  (state 2 single-state) | ASM**S**D**E**QAEARAFLSEEMIAEFKAAFDMFDADGGG**E**IS**A**KA**F**GTV**A**RM**NNVPVDPRVQEYVKRLT**D**Q**DGSGTI**S**FEEFLV**L**MV**KS**MK**Q**DA | 29 | 37 |
| High sequence identity state 2 single-state | ASMTDQQAEARAFLSEEMIAEFKAAFDMFDADGGGDISTKA**F**GTVMRMLG**V**NP**D**KE**VQ**D**Y**I**K**E**L**VD**Q**DGSGTIDFEEFLV**L**MVR**S**MKEDA | 11 | 25 |
| Multi-state designs (variable X) | ASM**E**D**L**QAEARAFLSEEMIAEFKAAFDMFDADGGGDIS**Y**KA**V**GTV**F**RMLG**I**NP**S**KE**V**LD**YLK**E**KI**D**V**DGSGTIDFEEFLV**L**MV**YX**MK**Q**DA | 18 | 25 |

**Table S2: Sequence differences between natural Ca^2+^-binding protein (1SMG) and computational designs.** Amino acid differences compared to 1SMG are red and bolded for the single-state state 2 design, the high sequence identity single-state state 2 design (as detailed in Fig. 1B), and the multi-state switch designs. Designable residues in multi-state design include all positions with different amino acids between states (red, bolded) and their neighbors as defined in the Methods section (blue, underlined). Allosteric position 89 is denoted as X in the multi-state designed sequence, which is variable in amino acid identity.

| Res | ∆∂Hn [ppm] | ∆∂N [ppm] | Res | ∆∂Hn [ppm] | ∆∂N [ppm] | Res | ∆∂Hn [ppm] | ∆∂N [ppm] | Res | ∆∂Hn [ppm] | ∆∂N [ppm] | Res | ∆∂Hn [ppm] | ∆∂N [ppm] |
| --- | --- | --- | --- | --- | --- | --- | --- | --- | --- | --- | --- | --- | --- | --- |
| 5 | 0.076 | 0.376 | 25 | 0.032 | 0.004 | 45 | 0.061 | 0.479 | **65** | 0.099 | 0.479 | 85 | 0.046 | 2.222 |
| 6 | 0.046 | 0.273 | 26 | 0.069 | 0.376 | 46 | 0.053 | 0.273 | **66** | 0.023 | 0.103 | 86 | 0.145 | 3.829 |
| 7 | 0.015 | 0.103 | 27 | 0.053 | 0.034 | 47 | 0.488 | 0.684 | **67** | 0.123 | 0.258 | 87 | 0.175 | 0.718 |
| 8 | 0.069 | 0.034 | 28 | 0.053 | 0.137 | 48 | 0.114 | 0.342 | **68** | 0.031 | 0.034 | 88 | 0.389 | 1.504 |
| 9 | 0.015 | 0.068 | 29 | 0.008 | 0.171 | 49 | 0.099 | 0.376 | **69** | 0.273 | 1.352 | **89 *** | 0.267 | 5.401 |
| 10 | 0.031 | 0.137 | 30 | 0.043 | 0.275 | 50 | 0.053 | 0.991 | **70** | 0.137 | 0.513 | 90 | 0.153 | 1.698 |
| 11 | 0.015 | 0.171 | 31 | 0.015 | 0.103 | 51 | 0.031 | 1.333 | **71** | 0.114 | 0.649 | 92 | 0.244 | 0.684 |
| 12 | 0.015 | 0.137 | 32 | 0.061 | 0.137 | 52 | 0.076 | 0.137 | **72** | 0.001 | 0.454 | 93 | 0.221 | 0.820 |
| 13 | 0.010 | 0.119 | 33 | 0.092 | 0.068 | **53** | 0.130 | 0.068 | **73** | 0.313 | 0.479 | 94 | 0.114 | 0.068 |
| 14 | 0.023 | 0.103 | 34 | 0.023 | 0.103 | **54** | 0.015 | 0.171 | **74** | 0.061 | 0.205 |  |  |  |
| 15 | 0.038 | 0.034 | 35 | 0.076 | 0.308 | **55** | 0.293 | 0.322 | **75** | 0.359 | 0.581 |  |  |  |
| 16 | 0.008 | 0.068 | 36 | 0.015 | 0.137 | **56** | 0.114 | 0.957 | **76** | 0.092 | 1.367 |  |  |  |
| 17 | 0.046 | 0.068 | 37 | 0.084 | 0.342 | **57** | - | - | 77 | 0.008 | 0.786 |  |  |  |
| 18 | 0.084 | 0.068 | 38 | 0.023 | 0.103 | **58** | 0.214 | 0.034 | 78 | 0.031 | 0.497 |  |  |  |
| 19 | 0.061 | 0.068 | 39 | 0.046 | 0.239 | **59** | 0.154 | 0.843 | 79 | 0.015 | 0.068 |  |  |  |
| 20 | 0.015 | 0.034 | 40 | 0.008 | 0.137 | **60** | 0.008 | 0.444 | 80 | 0.000 | 0.103 |  |  |  |
| 21 | 0.015 | 0.000 | 41 | 0.031 | 0.444 | **61** | 0.122 | 0.000 | 81 | 0.069 | 0.103 |  |  |  |
| 22 | 0.031 | 0.012 | 42 | 0.107 | 0.205 | **62** | 0.025 | 0.326 | 82 | 0.084 | 0.889 |  |  |  |
| 23 | 0.020 | 0.056 | 43 | 0.214 | 0.137 | **63** | 0.252 | 0.342 | 83 | 0.038 | 0.410 |  |  |  |
| 24 | 0.061 | 0.034 | 44 | 0.031 | 1.367 | **64** | 0.069 | 0.068 | 84 | 0.076 | 0.205 |  |  |  |

**Table S3: Chemical shift difference between design I89 and S89.** The chemical shift difference for each peak in the ^1^H,^15^N-HSQC spectra of I89 and S89. The reshaped region (residues 53-76) is in bold. Shift differences are colored by magnitude. There are significant shift differences not only in the near vicinity of the point mutation at position 89, but also throughout the reshaped region and in neighboring residues (43, 44, 47 on helix B). Position 57 is a proline.

| Res | R_1ρ_ [s^-1^] | Res | R_1ρ_ [s^-1^] | Res | R_1ρ_ [s^-1^] | Res | R_1ρ_ [s^-1^] | Res | R_1ρ_ [s^-1^] |
| --- | --- | --- | --- | --- | --- | --- | --- | --- | --- |
| 5 | 6.25±0.59 | 25 | 10.57±0.53 | 45 | 11.85±0.76 | **65** | 10.09±0.42 | 85 | 11.26±0.74 |
| 6 | 6.41±0.79 | 26 | 10.66±0.56 | 46 | 10.85±0.58 | **66** | 10.43±0.52 | 86 | 10.41±0.49 |
| 7 | 7.04±0.38 | 27 | 11.30±0.64 | 47 | 9.80±1.32 | **67** | 14.64±0.81 | 87 | 10.55±0.55 |
| 8 | 8.55±0.36 | 28 | 10.54±0.46 | 48 | 10.13±0.46 | **68** | 11.74±0.69 | 88 | 10.34±0.48 |
| 9 | 8.93±0.44 | 29 | 11.49±1.25 | 49 | 10.68±0.74 | **69** | - | 89 | 10.00±0.47 |
| 10 | 8.77±0.55 | 30 | 10.01±0.45 | 50 | 10.60±0.41 | **70** | 12.06±0.49 | 90 | 10.02±0.43 |
| 11 | 9.80±0.65 | 31 | 10.89±0.48 | 51 | 10.03±0.48 | **71** | 11.31±0.51 | 91 | 9.80±0.43 |
| 12 | 9.17±0.42 | 32 | 10.70±0.47 | 52 | 11.12±0.42 | **72** | 12.14±0.73 | 92 | 8.06±0.29 |
| 13 | 8.70±0.38 | 33 | 11.42±0.64 | **53** | 10.00±0.41 | **73** | 16.86±1.15 | 93 | 5.56±0.58 |
| 14 | 9.80±0.45 | 34 | 9.26±0.41 | **54** | 9.62±0.45 | **74** | 12.90±0.65 | 94 | 1.47±1.12 |
| 15 | 9.01±0.44 | 35 | 7.19±0.30 | **55** | 11.83±1.24 | **75** | 15.31±2.57 |  |  |
| 16 | 9.71±0.40 | 36 | 8.00±0.51 | **56** | 9.43±0.78 | **76** | 10.91±0.50 |  |  |
| 17 | 9.71±0.43 | 37 | 8.55±0.43 | **57** | - | 77 | 17.45±0.92 |  |  |
| 18 | 8.85±0.38 | 38 | 8.06±0.36 | **58** | 11.42±0.56 | 78 | 12.80±0.70 |  |  |
| 19 | 8.62±0.35 | 39 | 10.44±0.37 | **59** | 9.80±0.43 | 79 | 10.18±0.49 |  |  |
| 20 | 10.49±0.50 | 40 | 9.90±0.46 | **60** | 10.33±0.40 | 80 | 10.26±0.44 |  |  |
| 21 | 10.70±0.48 | 41 | 18.32±1.16 | **61** | 10.45±0.49 | 81 | 10.65±0.50 |  |  |
| 22 | 10.60±0.48 | 42 | 11.24±0.50 | **62** | 10.43±0.48 | 82 | 10.47±0.49 |  |  |
| 23 | 9.80±0.43 | 43 | 14.56±0.85 | **63** | 10.70±0.61 | 83 | 10.11±0.49 |  |  |
| 24 | 10.82±0.50 | 44 | 14.95±0.80 | **64** | 10.56±0.46 | 84 | 10.48±0.53 |  |  |

**Table S4: R_1ρ_ values for apo I89.** The R_1ρ_ rate for each residue of apo design I89 colored by magnitude. The reshaped region (residues 53-76) is in bold. There are elevated relaxation rates throughout the reshaped region where the largest conformational change is expected (residues 66-75) and in residues facing that region (41, 42, 43, 44 in helix B). Position 57 is a proline. No value could be calculated for position 69 due to low intensity and peak overlap.

| Res | ∆∂Hn [ppm] | ∆∂N [ppm] | Res | ∆∂Hn [ppm] | ∆∂N [ppm] | Res | ∆∂Hn [ppm] | ∆∂N [ppm] | Res | ∆∂Hn [ppm] | ∆∂N [ppm] | Res | ∆∂Hn [ppm] | ∆∂N [ppm] |
| --- | --- | --- | --- | --- | --- | --- | --- | --- | --- | --- | --- | --- | --- | --- |
| 5 | 0.049 | 0.256 | 25 | 0.027 | 0.053 | 45 | 0.067 | 0.934 | **65** | 0.012 | 0.364 | 85 | 0.098 | 0.787 |
| 6 | 0.053 | 0.376 | 26 | 0.009 | 0.019 | 46 | 0.124 | 0.459 | **66** | 0.226 | 0.828 | 86 | 0.080 | 0.188 |
| 7 | 0.078 | 0.036 | 27 | 0.013 | 0.252 | 47 | 0.260 | 0.195 | **67** | 0.136 | 1.639 | 87 | 0.048 | 0.089 |
| 8 | 0.038 | 0.084 | 28 | 0.004 | 0.153 | 48 | 0.028 | 0.038 | **68** | 0.171 | 0.892 | 88 | 0.001 | 0.182 |
| 9 | 0.010 | 0.350 | 29 | 0.067 | 0.192 | 49 | 0.129 | 0.045 | **69** | - | - | 89 | 0.078 | 0.116 |
| 10 | 0.081 | 0.336 | 30 | 0.071 | 0.125 | 50 | 0.026 | 0.259 | **70** | 0.063 | 0.572 | 90 | 0.034 | 0.103 |
| 11 | 0.104 | 0.030 | 31 | 0.009 | 0.257 | 51 | 0.003 | 0.009 | **71** | 0.203 | 2.017 | 91 | 0.037 | 0.073 |
| 12 | 0.042 | 0.194 | 32 | 0.009 | 0.157 | 52 | 0.053 | 0.117 | **72** | 0.002 | 1.395 | 92 | 0.001 | 0.132 |
| 13 | 0.023 | 0.173 | 33 | 0.018 | 0.113 | **53** | 0.031 | 0.174 | **73** | 0.204 | 0.987 | 93 | 0.004 | 0.105 |
| 14 | 0.004 | 0.196 | 34 | 0.110 | 0.004 | **54** | 0.020 | 0.201 | **74** | 0.053 | 0.221 | 94 | 0.007 | 0.045 |
| 15 | 0.027 | 0.098 | 35 | 0.019 | 0.561 | **55** | 0.075 | 0.419 | **75** | - | - |  |  |  |
| 16 | 0.001 | 0.040 | 36 | 0.061 | 0.893 | **56** | 0.001 | 0.160 | **76** | 0.082 | 1.293 |  |  |  |
| 17 | 0.004 | 0.057 | 37 | 0.149 | 0.742 | **57** | - | - | 77 | - | - |  |  |  |
| 18 | 0.029 | 0.012 | 38 | 0.124 | 0.251 | **58** | 0.231 | 0.830 | 78 | 0.203 | 1.231 |  |  |  |
| 19 | 0.001 | 0.004 | 39 | 0.083 | 0.058 | **59** | 0.053 | 0.127 | 79 | 0.096 | 0.213 |  |  |  |
| 20 | 0.005 | 0.020 | 40 | 0.062 | 0.692 | **60** | 0.061 | 0.295 | 80 | 0.009 | 0.223 |  |  |  |
| 21 | 0.009 | 0.008 | 41 | 0.207 | 1.792 | **61** | 0.062 | 0.085 | 81 | 0.036 | 0.081 |  |  |  |
| 22 | 0.010 | 0.098 | 42 | 0.001 | 0.613 | **62** | 0.097 | 0.230 | 82 | 0.098 | 0.574 |  |  |  |
| 23 | 0.027 | 0.184 | 43 | - | - | **63** | 0.161 | 0.219 | 83 | 0.019 | 0.076 |  |  |  |
| 24 | 0.012 | 0.119 | 44 | 0.006 | 0.973 | **64** | 0.033 | 0.336 | 84 | 0.026 | 0.111 |  |  |  |

**Table S5: Chemical shift perturbations for I89 with the addition of Ca^2+^.** The chemical shift difference for each peak in the ^1^H,^15^N-HSQC spectra of I89 with and without 10eq. of Ca^2+^ added. The reshaped region (residues 53-76) is in bold. Ca^2+^ binding sites I and II are underlined. Shift differences are colored by magnitude. There are significant shift differences not only in the Ca^2+^ binding sites (site I: residues 34-40, site II: residues 70-76 and 81), but also in distal areas throughout the reshaped region and in residues facing the reshaped region from neighboring helices (44, 47 on helix B; 6, 10, 11 on helix N). Peak 57 is a proline. For other missing values (i.e. residues 43, 69, 75, 77), an assignment could not be made for S89 and/or I89 with Ca^2+^ added.

|  | **I89** | | **S89** | |
| --- | --- | --- | --- | --- |
|  | **Ca^2+^-free** | **+Ca^2+^** | **Ca^2+^-free** | **+Ca^2+^** |
| **Box dimensions (nm)** | 3.569 x 3.399 x 4.134 nm | 3.569 x 3.399 x 4.134 nm | 3.790 x 4.029 x 3.706 nm | 3.790 x 4.029 x 3.706 nm |
| **K^+^ ions** | 22 | 22 | 23 | 23 |
| **Cl^-^ ions** | 11 | 13 | 12 | 14 |
| **Ca^2+^ ions** | 0 | 1 | 0 | 1 |
| **Water molecules** | 5718 | 5715 | 5983 | 5980 |

**Table S6: Molecular dynamics simulation details.** Solvation box details for I89 and S89 +/- Ca^2+^ MD trajectories.

| **Structure** | **Cα RMSD to state 1**  **Reshaped helix (whole ordered structure) [Å]** | **Cα RMSD to state 2**  **Reshaped helix (whole ordered structure) [Å]** |
| --- | --- | --- |
| Single-state #6306 (NMR) | 4.99 (2.38) | 1.22 (1.21) |
| S89 (NMR) | 5.04 (2.42) | 1.33 (1.31) |
| I89 *apo* proposal #1 (NMR) | 2.00 (1.67) | 4.85 (2.39) |
| I89 *apo* proposal #2 (NMR) | 5.04 (2.42) | 1.33 (1.31) |
| I89 +Ca^2+^ (NMR) | 1.18 (1.34) | 5.49 (2.53) |

**Table S7: The Cα RMSDs to states 1 and 2 of each experimentally solved structure.** The Cα RMSD values for the reshaped helix (residues 59-69) and the whole structure (excluding loops) for the experimentally determined structures of each design align well with either the top-ranking (by average pLDDT) AF2 model for I89 (state 1) or the top-ranking AF2 model for S89 (state 2) as expected (green shading). The side chain-level details for each structure and their corresponding AF2 model can be found in Fig. S5.

| **Design ID** | **DNA sequence** | **Protein sequence** |
| --- | --- | --- |
| 415 | ggagggtcggcttcgcatatgCATCACCATCACCATCACAGCAGCGGCGCTGTGCCGGAAGGCAGCCATGCTGCCAGCATGGACGACGAACAAGCAGAAGCAAGGGCCTTCCTTTCTGAAGAGATGATTGCAGAATTTAAAGCCGCTTTCGATATGTTTGATGCTGATGGCGGTGGGGAAATATCGATAAAAGCGGCTGGAACATTATCAAGAATGATTAACGCTCCACCTACCGAGGAGGTCTTGCAGAAAATTCTAAAGAGAAAGGATAATGATGGTAGTGGTACTATCTCCTTTGAAGAATTTCTGGTAGCAATGGTTTACTATATGAAGGATGACGCTctcgagggtggaggttccgaacaacagcttatttctgaagaggacttgta | ASMDDEQAEARAFLSEEMIAEFKAAFDMFDADGGGEISIKAAGTLSRMINAPPTEEVLQKILKRKDNDGSGTISFEEFLVAMVYYMKDDA |
| 567 | ggagggtcggcttcgcatatgCATCACCATCACCATCACAGCAGCGGCGCTGTGCCGGAAGGCAGCCATGCTGCTTCTATGTCAGATGAACAAGCTGAGGCAAGAGCCTTTTTGTCCGAGGAAATGATAGCAGAGTTTAAGGCAGCGTTCGACATGTTTGATGCCGACGGGGGTGGCGAAATAAGTGCCAAAGCTGCGGGTACGATTATTAGGATGCTTAATCTGGATGAAACTGCTGCAAAATGGATTAAAAAAAAGGCAGAAAAAGACACAGATGGTTCGGGAACCATCTCTTTCGAAGAATTTTTACTAGCTATGGTCGTTGTAATGAAGAGCGATGCTctcgagggtggaggttccgaacaacagcttatttctgaagaggacttgta | ASMSDEQAEARAFLSEEMIAEFKAAFDMFDADGGGEISAKAAGTIIRMLNLDETAAKWIKKKAEKDTDGSGTISFEEFLLAMVVVMKSDA |
| 615 | ggagggtcggcttcgcatatgCATCACCATCACCATCACAGCAGCGGCGCTGTGCCGGAAGGCAGCCATGCTGCCGTGATGACTGACGAAGCAGCAGAAGCTAGGGCTTTTCTTAGCGAAGAAATGATTGCTGAGTTTAAAGCTGCATTCGATATGTTTGATGCGGATGGTGGGGGCTCAATCAGTAAGAAAGCGCTAGGAACGTACCAAAGAATGACTGGTGTACCATTCGATCCTAACAAGGCTGAGTCTTTAGCCAGACGTTATGATAATGATGGTTCCGGAACCGTCACATTTGACGAATTCTTGATAGTTATGGTTCAGGACATGAAAGCCGATGCActcgagggtggaggttccgaacaacagcttatttctgaagaggacttgta | AVMTDEAAEARAFLSEEMIAEFKAAFDMFDADGGGSISKKALGTYQRMTGVPFDPNKAESLARRYDNDGSGTVTFDEFLIVMVQDMKADA |
| 1154 | ggagggtcggcttcgcatatgCATCACCATCACCATCACAGCAGCGGCGCTGTGCCGGAAGGCAGCCATGCTGCCAAAATGGACGATCTACAAGCCGAAGCAAGAGCATTTTTATCCGAAGAGATGATCGCCGAATTTAAAGCTGCTTTTGACATGTTTGACGCGGATGGTGGTGGTAGAATTAGTCCTAAAGCTCTCGGGACCATTGCAAGGATGACAAATGTAGCTGATCCAAAAGAATTGAAGAAGGCTGCAAAGTATATACAGGATCGTGATGGATCAGGCACGTTAACTTTCGAGGAATTCCTTCTGGCTATGGTTTTGGTCATGAAATCTGATGCGctcgagggtggaggttccgaacaacagcttatttctgaagaggacttgta | AKMDDLQAEARAFLSEEMIAEFKAAFDMFDADGGGRISPKALGTIARMTNVADPKELKKAAKYIQDRDGSGTLTFEEFLLAMVLVMKSDA |
| 2114 | ggagggtcggcttcgcatatgCATCACCATCACCATCACAGCAGCGGCGCTGTGCCGGAAGGCAGCCATGCTGCGAGTATGGATGATGAGCAAGCTGAAGCTAGGGCATTCTTGTCAGAAGAAATGATTGCCGAATTTAAAGCAGCTTTTGACATGTTCGATGCAGATGGTGGTGGTGAGATTTCCGCCAAAGCACTGGGAACTTGGGCTAGAATGCTTAAACAGATCAATCCAGCTGTAGAAAAGGCTGCAAAGAAGGCCAAAGACGATGCGGATGGGTCTGGCACAATAGAATTTGAAGAGTTCCTAAAGTTAATGGTTCAATATATGAAGAAAGACGCCctcgagggtggaggttccgaacaacagcttatttctgaagaggacttgta | ASMDDEQAEARAFLSEEMIAEFKAAFDMFDADGGGEISAKALGTWARMLKQINPAVEKAAKKAKDDADGSGTIEFEEFLKLMVQYMKKDA |
| 3880 | ggagggtcggcttcgcatatgCATCACCATCACCATCACAGCAGCGGCGCTGTGCCGGAAGGCAGCCATGCTGCAAGCATGGATGACAAACAGGCTGAAGCTAGAGCATTTCTATCTGAGGAAATGATAGCCGAATTCAAGGCTGCGTTTGATATGTTTGATGCCGATGGAGGTGGTGAAATCTCCCCAAAAGCCGCAGGTACAGTATTTAGGATGCAAAATTTAGATGAAAAGAAAGCTAGAGAAGCTGTTGAGGAGTGGAAAAAGGATAAAGACGGCTCAGGGACTATTAGTTTCGAAGAATTCTTGATAATTATGGTCTGGATTATGAAGGACGATGCActcgagggtggaggttccgaacaacagcttatttctgaagaggacttgta | ASMDDKQAEARAFLSEEMIAEFKAAFDMFDADGGGEISPKAAGTVFRMQNLDEKKAREAVEEWKKDKDGSGTISFEEFLIIMVWIMKDDA |
| 4602 | ggagggtcggcttcgcatatgCATCACCATCACCATCACAGCAGCGGCGCTGTGCCGGAAGGCAGCCATGCTGCAAAGATGGACGATTTAGTAGCTGAGGCACGTGCTTTTCTGTCTGAGGAAATGATAGCCGAATTTAAAGCGGCATTTGATATGTTTGACGCCGATGGTGGTGGTAAAATCTCAGCAAAAGCTGCAGGGACATTCGCTAGAATGCTTAATTTGGATCCAAGGAAGTTCGAACGAGCTGCCAGGGAAATTGCTAGAGATGTTGACGGAAGTGGCACTGTTTCCTTTGAAGAATTCCTAGCGGCCATGGTCGCTGTGATGAAGAGAGATGCActcgagggtggaggttccgaacaacagcttatttctgaagaggacttgta | AKMDDLVAEARAFLSEEMIAEFKAAFDMFDADGGGKISAKAAGTFARMLNLDPRKFERAAREIARDVDGSGTVSFEEFLAAMVAVMKRDA |
| 4851 | ggagggtcggcttcgcatatgCATCACCATCACCATCACAGCAGCGGCGCTGTGCCGGAAGGCAGCCATGCTGCTTCGATGGATGATGAACAAGCAGAAGCAAGAGCATTTTTAAGCGAAGAGATGATTGCCGAATTTAAAGCCGCTTTTGACATGTTCGACGCGGACGGTGGTGGTTCTATATCCGCTAAAGCATTAGGCACTGCTGCTAGGATGCTAAATCTTGATGAGGAAGCCGCTAAGAAATGGGCCAAGAAAGCGCAGGACGATCTGGATGGGTCAGGAACAGTAAGTTTCGAAGAGTTTTTGTTGTGGATGGTTTGGGTCATGAAGGATGATGCActcgagggtggaggttccgaacaacagcttatttctgaagaggacttgta | ASMDDEQAEARAFLSEEMIAEFKAAFDMFDADGGGSISAKALGTAARMLNLDEEAAKKWAKKAQDDLDGSGTVSFEEFLLWMVWVMKDDA |
| 5505 | ggagggtcggcttcgcatatgCATCACCATCACCATCACAGCAGCGGCGCTGTGCCGGAAGGCAGCCATGCTGCTAGTATGGACGATGAACAAGCAGAAGCAAGAGCTTTTTTGTCCGAGGAAATGATCGCAGAATTTAAGGCGGCATTCGATATGTTTGACGCCGATGGCGGGGGTGGTATTTCTCCAAAAGCGGCTGGAACAATTGCCAGGATGTTAAATCTTGACGAAGACAAAGCCAGAAAGGTTGCTAAAAAGTTGTTAGATGATAAAGATGGAAGCGGTACTATATCATTCGAGGAATTTCTAGTAGTCATGGTGATAATTATGAAAGATGATGCTctcgagggtggaggttccgaacaacagcttatttctgaagaggacttgta | ASMDDEQAEARAFLSEEMIAEFKAAFDMFDADGGGGISPKAAGTIARMLNLDEDKARKVAKKLLDDKDGSGTISFEEFLVVMVIIMKDDA |
| 6306 | ggagggtcggcttcgcatatgCATCACCATCACCATCACAGCAGCGGCGCTGTGCCGGAAGGCAGCCATGCTGCTTCTATGAGTGATGAACAAGCTGAAGCTAGGGCATTCTTGAGCGAAGAGATGATTGCTGAGTTTAAGGCAGCCTTCGACATGTTTGATGCAGATGGTGGTGGCGAAATATCCGCTAAAGCGTTTGGTACCGTCGCCCGTATGAACAATGTTCCAGTGGACCCTAGAGTCCAAGAATATGTAAAAAGACTTACAGACCAAGATGGATCTGGGACTATTTCATTTGAGGAATTCTTAGTTCTAATGGTTAAATCAATGAAGCAGGATGCCctcgagggtggaggttccgaacaacagcttatttctgaagaggacttgta | ASMSDEQAEARAFLSEEMIAEFKAAFDMFDADGGGEISAKAFGTVARMNNVPVDPRVQEYVKRLTDQDGSGTISFEEFLVLMVKSMKQDA |
| 7246 | ggagggtcggcttcgcatatgCATCACCATCACCATCACAGCAGCGGCGCTGTGCCGGAAGGCAGCCATGCTGCAAGTATGGACGATAGACAAGCCGAAGCAAGGGCTTTCCTTTCAGAAGAGATGATTGCTGAATTCAAGGCTGCCTTTGACATGTTTGACGCTGATGGTGGGGGTGAAATTTCAGCTAAAGCGGCCGGAACTGTTTTGAGAATGGCAAATGTCCCAGCCGATGATGAGATTAAGGAAGAAATAAAAAAGTTAGCTGACGATGATGGTTCCGGCACAATCGATTTTGAGGAATTTCTACGTCTGATGGTAAAAGCAATGAAATCTGATGCActcgagggtggaggttccgaacaacagcttatttctgaagaggacttgta | ASMDDRQAEARAFLSEEMIAEFKAAFDMFDADGGGEISAKAAGTVLRMANVPADDEIKEEIKKLADDDGSGTIDFEEFLRLMVKAMKSDA |
| 1smg WT | ggagggtcggcttcgcatatgCATCACCATCACCATCACAGCAGCGGCGCTGTGCCGGAAGGCAGCCATGCTGCTAGTATGACAGATCAACAAGCTGAAGCAAGAGCCTTCCTATCCGAGGAAATGATTGCCGAATTTAAGGCTGCATTTGATATGTTCGACGCGGACGGTGGTGGCGATATATCTACGAAGGCACTGGGAACTGTCATGAGAATGTTAGGACAGAATCCAACTAAAGAAGAATTGGATGCTATCATTGAAGAGGTGGATGAGGATGGCTCAGGTACCATAGACTTTGAAGAGTTTCTTGTAATGATGGTTAGGCAGATGAAAGAAGATGCCctcgagggtggaggttccgaacaacagcttatttctgaagaggacttgta | ASMTDQQAEARAFLSEEMIAEFKAAFDMFDADGGGDISTKALGTVMRMLGQNPTKEELDAIIEEVDEDGSGTIDFEEFLVMMVRQMKEDA |

**Table S8. DNA and protein sequences of single state designs selected for experimental testing.** The coding portion of the DNA is in uppercase. Homology regions used for yeast transformation are in lowercase.

| **Design ID** | **DNA sequence** | **Protein sequence** |
| --- | --- | --- |
| I89 | GCGTCGATGGAGGATTTGCAAGCCGAGGCTCGGGCCTTTCTGTCAGAGGAGATGATTGCCGAGTTTAAGGCCGCGTTTGATATGTTTGATGCCGATGGTGGCGGGGATATATCTTACAAAGCCGTAGGGACCGTATTTCGTATGCTGGGTATCAATCCATCCAAGGAAGTCTTGGATTATCTGAAAGAAAAAATTGATGTAGACGGTTCAGGCACAATTGATTTCGAAGAATTTCTGGTTTTGATGGTATACATCATGAAACAGGACGCATAActcgagcctgatgcggtattttctcct | ASMEDLQAEARAFLSEEMIAEFKAAFDMFDADGGGDISYKAVGTVFRMLGINPSKEVLDYLKEKIDVDGSGTIDFEEFLVLMVYIMKQDA |
| K89 | GCCTCAATGGAGGATCTTCAAGCAGAGGCACGAGCGTTTTTAAGTGAGGAGATGATCGCCGAATTTAAAGCGGCATTCGACATGTTTGACGCAGACGGTGGTGGCGATATCAGTTACAAAGCCGTCGGTACCGTATTTCGTATGTTGGGGATAAACCCGTCGAAGGAAGTTCTGGATTACCTTAAAGAAAAAATTGATGTGGATGGAAGTGGTACTATTGATTTTGAGGAGTTTTTGGTGCTTATGGTGTATAAGATGAAGCAAGATGCCTGActcgagcctgatgcggtattttctcct | ASMEDLQAEARAFLSEEMIAEFKAAFDMFDADGGGDISYKAVGTVFRMLGINPSKEVLDYLKEKIDVDGSGTIDFEEFLVLMVYKMKQDA |
| N89 | GCATCTATGGAGGACCTGCAGGCCGAGGCTAGAGCTTTTCTCAGTGAAGAAATGATAGCCGAATTTAAAGCTGCATTTGATATGTTCGATGCCGATGGTGGCGGTGATATTAGCTATAAAGCGGTTGGTACTGTGTTTCGCATGCTTGGCATTAACCCATCAAAGGAAGTCCTGGACTATTTGAAAGAAAAAATCGATGTCGATGGCAGTGGTACCATTGATTTTGAGGAATTTCTGGTTCTTATGGTATACAATATGAAGCAGGATGCATAActcgagcctgatgcggtattttctcct | ASMEDLQAEARAFLSEEMIAEFKAAFDMFDADGGGDISYKAVGTVFRMLGINPSKEVLDYLKEKIDVDGSGTIDFEEFLVLMVYNMKQDA |
| S89 | GCCAGTATGGAGGATTTGCAGGCTGAAGCCAGAGCATTTTTGTCAGAAGAAATGATAGCCGAGTTTAAGGCCGCGTTCGACATGTTTGATGCCGACGGTGGGGGCGATATTTCCTATAAAGCAGTCGGTACAGTATTCCGTATGCTGGGTATCAATCCATCAAAGGAAGTTCTCGACTATCTGAAAGAAAAAATTGACGTTGATGGTAGTGGGACAATTGATTTTGAGGAGTTCTTAGTATTAATGGTTTATTCCATGAAACAGGATGCTTAActcgagcctgatgcggtattttctcct | ASMEDLQAEARAFLSEEMIAEFKAAFDMFDADGGGDISYKAVGTVFRMLGINPSKEVLDYLKEKIDVDGSGTIDFEEFLVLMVYSMKQDA |
| R89 | GCATCGATGGAAGACCTGCAAGCGGAAGCGCGTGCATTCCTGAGCGAGGAAATGATTGCCGAGTTTAAAGCTGCCTTCGACATGTTTGATGCCGACGGAGGAGGTGATATTTCATATAAGGCTGTGGGTACTGTATTTCGGATGCTCGGGATTAATCCTAGTAAAGAAGTATTGGACTACCTTAAAGAAAAAATCGACGTGGATGGGTCCGGGACTATAGATTTTGAAGAATTTCTCGTTCTGATGGTCTATCGGATGAAACAAGATGCCTAGctcgagcctgatgcggtattttctcct | ASMEDLQAEARAFLSEEMIAEFKAAFDMFDADGGGDISYKAVGTVFRMLGINPSKEVLDYLKEKIDVDGSGTIDFEEFLVLMVYRMKQDA |
| I89 Y64F | GCCTCGATGGAGGATTTGCAAGCGGAGGCCCGCGCTTTCCTTTCGGAGGAAATGATTGCGGAATTTAAAGCCGCATTCGATATGTTTGACGCTGATGGTGGGGGCGATATTAGCTATAAAGCCGTAGGTACTGTGTTCAGAATGCTTGGTATTAACCCCTCAAAAGAAGTTCTGGACTTTCTTAAGGAAAAAATCGACGTGGACGGAAGCGGTACGATCGATTTTGAAGAATTTCTGGTATTGATGGTATACATTATGAAGCAGGATGCCTAActcgagcctgatgcggtattttctcct | ASMEDLQAEARAFLSEEMIAEFKAAFDMFDADGGGDISYKAVGTVFRMLGINPSKEVLDFLKEKIDVDGSGTIDFEEFLVLMVYIMKQDA |
| I89 K68E | GCATCTATGGAAGACCTGCAGGCTGAGGCACGGGCCTTTTTATCTGAAGAAATGATCGCAGAATTTAAAGCCGCGTTTGATATGTTCGATGCAGATGGCGGTGGCGATATTTCCTATAAGGCGGTTGGTACGGTTTTTCGTATGTTGGGGATTAATCCCTCAAAAGAAGTACTGGATTATCTTAAGGAGGAGATCGATGTTGACGGAAGTGGAACCATCGACTTTGAGGAATTTCTCGTCCTTATGGTGTACATTATGAAGCAGGATGCTTAActcgagcctgatgcggtattttctcct | ASMEDLQAEARAFLSEEMIAEFKAAFDMFDADGGGDISYKAVGTVFRMLGINPSKEVLDYLKEEIDVDGSGTIDFEEFLVLMVYIMKQDA |

**Table S9. DNA and protein sequences of multi-state designs selected for experimental testing.** The coding portion of the DNA sequence is in uppercase.

|  | **Single-state design #6306** | **Switch design S89** | **Switch design I89 *apo* state 1** | **Switch design I89 *apo* state 2** | **Switch design I89 +Ca^2+^** |
| --- | --- | --- | --- | --- | --- |
| Number of residues | 94 | 94 | 94 | 94 | 94 |
| Total number of structures computed | 100 | 100 | 100 | 100 | 100 |
| Number of structures reported | 20 | 20 | 20 | 20 | 20 |
| Backbone atoms RMSD (Å) | 0.92 | 0.96 | 0.47 | 0.60 | 0.57 |
| Heavy atoms RMSD (Å) | 1.31 | 1.35 | 0.83 | 0.91 | 0.90 |
| **Restraints** | | | | |  |
| Total distance restraints | 2052 | 1820 | 2498 | 2393 | 2285 |
| Intra-residue [ i = j ] | 499 | 425 | 583 | 585 | 496 |
| Sequential [ \| i - j \| = 1 ] | 509 | 467 | 591 | 572 | 604 |
| Medium-range [ 1 < \| i - j \| < 5 ] | 528 | 495 | 677 | 610 | 612 |
| Long range [ \| i - j \| > 4 ] | 516 | 433 | 647 | 626 | 573 |
| Dihedral angle restraints | 156 | 164 | 168 | 168 | 170 |
| **Violations** | | | | |  |
| Ave. distance restraints (Å) | 0.19±0.08 | 0.2±0.1 | 0.2±0.1 | 0.18±0.09 | 0.18±0.08 |
| Ave. dihedral angle restraints (°) | 1.3±0.5 | 3.1±2.1 | 3.8±3.0 | 3.3±3.1 | 3.3±2.9 |
| Max. distance restraint violation (Å) | 0.59 | 0.73 | 0.74 | 0.92 | 0.56 |
| Max. dihedral angle restraint violation (°) | 4.66 | 10.16 | 12.91 | 13.79 | 14.75 |
| Num. distance violations > 0.2Å | 4.4 | 4.8 | 11.9 | 6.4 | 11.1 |
| Num. dihedral angle violations > 10° | 0.0 | 0.1 | 1.2 | 1.2 | 1.6 |
| **Model quality** | | | | |  |
| *Deviations from idealized geometry* | | | | |  |
| Bond lengths (Å) | None | None | None | None | None |
| Bond angles (°) | None | None | None | None | None |
| *Ramachandran* | | | | |  |
| Favored regions | 96±2% | 98±1% | 97±1% | 96±1% | 96±1% |
| Allowed regions | 3±2% | 2±1% | 3±1% | 4±1% | 4±1% |
| Disallowed regions | 0±0% | 0±0% | 0±0% | 0±0% | 0±0% |
| Clash score | 6 | 5 | 8 | 7 | 10 |

**Table S10: NMR structural statistics.**
